# Inferential reasoning in non-humans: a critical analysis of experimental evidence

**DOI:** 10.1101/2025.09.23.678036

**Authors:** Dániel Rivas-Blanco, Marco Vasconcelos, Friederike Range, Alex Kacelnik, Tiago Monteiro

**Affiliations:** Domestication Lab, Konrad Lorenz Institute of Ethology, Department of Interdisciplinary Life Sciences, University of Veterinary Medicine Vienna; Vienna, 1210 Vienna, Austria; William James Center for Research, University of Aveiro; Aveiro, 3810-193 Aveiro, Portugal; Department of Biology, University of Oxford, Oxford, OX1 3PS Oxford, UK

## Abstract

Is human language necessary for logical reasoning? Multiple studies address this by showing animals a bait hidden in either of two inverted cups, briefly lifting one exposing its content and then letting subjects choose. Consistent preference for the baited container is interpreted as evidence of inferential reasoning by exclusion (“*if the bait is not here, it must be there*”). More complex variations invoke metacognitive revision of hypotheses. We show that these protocols do not demonstrate logical reasoning. Instead, each event-stimulus elicits different actions and this suffices to choose correctly between them. We demonstrate this with experimental starlings and argue that other species may show similar behaviour, highlighting the need for careful controls when attributing high-level mental capacities such as reasoning or causal understanding.

## Introduction

### Demonstrating inferential reasoning

The quest to infer mental events from observable behaviour is a persistent source of both inspiration and tension for different research programmes. Comparative cognition research benefits from a rich theoretical contrast between schools that avoid reliance on mental assumptions, such as Skinner’s radical behaviourism (see 1), and at the other extreme those that invoke high-level constructs such as ‘causal understanding’ as explanatory mechanisms for animal behaviour (e.g. 2). Following Heyes (3), we refer to the use of high-level cognitive constructs as explanatory tools as super-cognitivism, to highlight the contrast with both radical behaviourism and cognitive approaches that model psychological processes algorithmically, using observable behaviour as both input and output. Scholars adept to either radical behaviourism or supercognitivism often respectively criticise perceived excesses in anthropomorphism (4) or anthropodenialism (5), perhaps with some justification.

We argue theoretically and show experimentally that some observations widely attributed to high-level constructs such as logical reasoning and/or causal understanding can be modelled precisely and productively in terms of observable behaviour, including cases in which the data have been interpreted as justifying supercognitive claims. We focus on the *2-Cups Task* paradigm (**Figure 1A**) (6,7) as described in detail in a widely subscribed proposal by the *ManyPrimates* research consortium (8). We aim to show that an algorithmic model that is simple for agents to perform, empirically verifiable, and frugal in parameters can (and does) generate flexible behaviour in that and multiple alternative scenarios. Our proposal aims to implement objectives expressed in behavioural ecology (9), in comparative cognition (10) and in human decision research (11). Shettleworth (10) expressed the resistance to account for interesting complex behaviour through simple algorithms succinctly:

**Figure 1.**
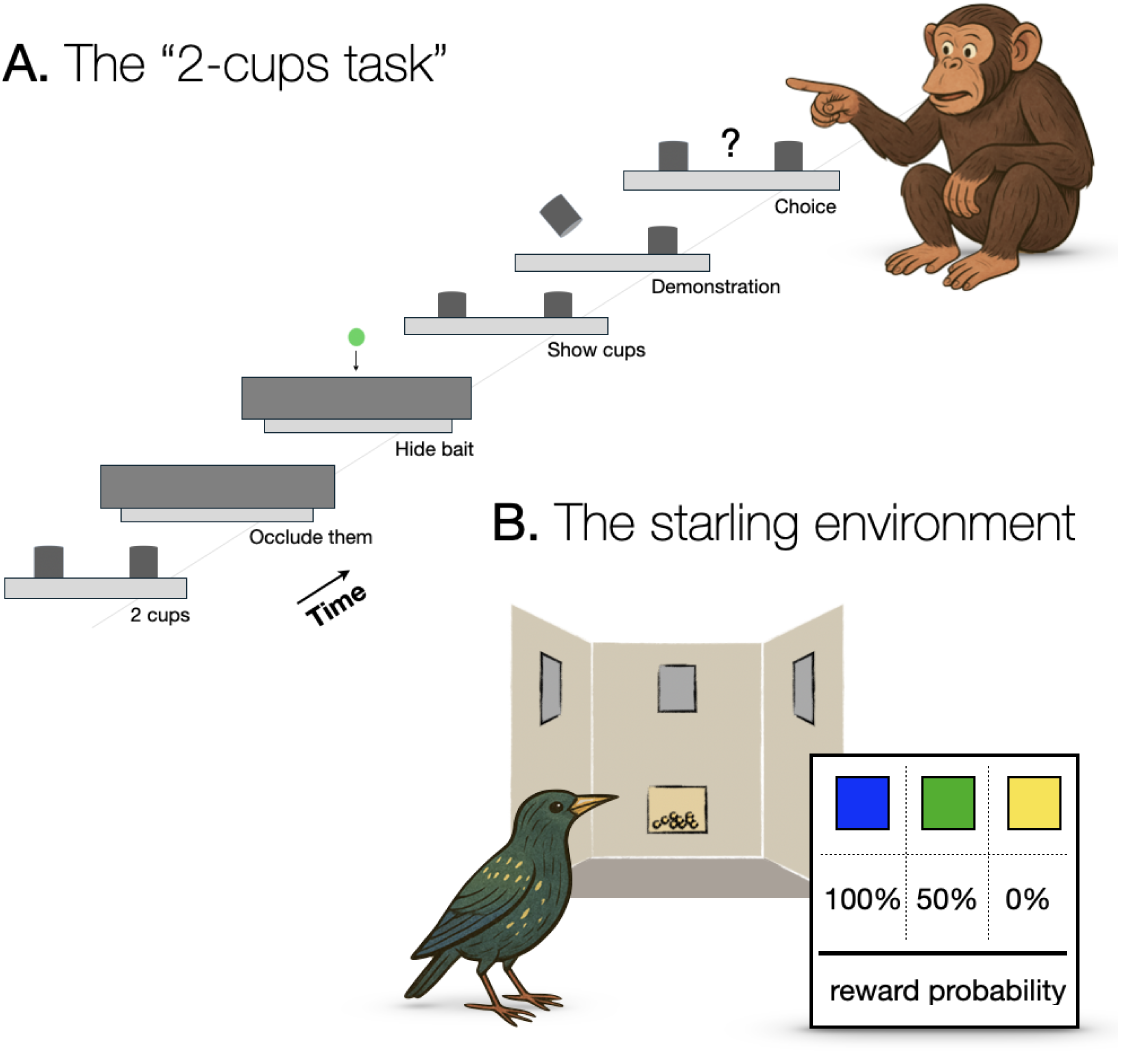
The 2-cups task. **(A**) Example trial in the 2-cups protocol where the subject is shown the content of the empty cup before being allowed to make a choice. (**B**) Emulation of the 2-cups task with starlings using a 3-key operant panel and 3 coloured stimuli signalling *Sure Reward* (100%), sure *No Reward* (0%), or *Uncertain* (50%). Animal images were generated using ChatGPT (https://chat.openai.com).

*“…demonstrations that complex human-like behavior arises from simple mechanisms rather than from ‘higher’ processes, such as insight or theory of mind, are often seen as uninteresting and ‘killjoy’, almost a denial of mental continuity between other species and humans.*”

Within this spirit, we propose a simple mechanism that can account for complex adaptive behaviour in many different situations, including those supporting the attribution of inferential logic to apes.

The *ManyPrimates 3A - Inference by Exclusion* proposal starts from the premise that individuals in nature need to make inferences about hidden information (8). We quote:

*“Important events are often not observable. Consequently, individuals must make inferences about unobserved events. One method for making such inferences is inference by exclusion (IbE) - the ability to reconstruct the identity of an unknown alternative by eliminating known options.”*

We believe that this premise is theoretically misleading and examine it experimentally. A useful theoretical starting point is Gibson’s (12) concept of affordances in visual perception. He argued that evolution shapes organisms to perceive and act upon what matters to them in their environment, writing that *to perceive visual stimuli is to perceive what they afford*. For him perception is about enabling action, not representing or ‘understanding’ the world. Gibson’s ecological view of perception needs not be universally shared, but for present purposes we argue that if an event is important necessarily it must have detectable consequences and species would have evolved mechanisms to act adaptively towards them without necessarily making mental inferences about what they cannot directly see or understand.

The often claimed evidence for inferential reasoning downplays the potential of simple learning processes evolutionarily designed to use reliable predictors to generate adaptive behaviour. To avoid confusions, we note that some authors use the phrase ‘Inference by Exclusion’ to address observable behaviour rather than mental processes (see discussion by 13), but here we follow its most widespread and conceptually ambitious use, in which making inferences and following logical reasoning ‘explains’ behaviour. The supercognitivist handling of *Inference by Exclusion* does not just include logical operations, but implicitly involves psychological object permanence, representation of probabilities and causal understanding. We will argue that none of these is required to account for the reported data. To support our claims, we apply an alternative choice model to an experimental emulation of the *2-Cups Task*, using spotless starlings (*Sturnus unicolor*) as subjects (**Figure 1B**) and suggest that our alternative framework may be useful for primates and other taxa. We do not predicate the absence of reasoning processes, but object to some claims for evidence of their presence and show that there exist approaches to modelling cognition that are epistemically preferable for their frugality, ability to generate testable predictions and greater vulnerability to empirical falsification. We describe our model in the next section, then examine the *2-Cups Task* in some detail and present our experimental emulation.

### Logical reasoning or independent responding

When a goal-directed agent faces mutually exclusive options, it is tempting to model the agent as if it reasoned about the options’ relative merits with respect to its goal, and then made a choice determined by these mental deliberations. This implies that preference is constructed when facing the choice. This approach fits human subjective conscious experience and dominates the field of human decision making. However, the agent can be alternatively conceptualised as possessing a pattern of behaviour towards each option on its own that is also elicited when facing more than one option simultaneously. In a choice the candidate behaviours compete for a mutually exclusive *Behavioural Final Common Path* (14), so that only one action is realised. The implication is that preference expresses cognitive contents that exist before the subject faces the choice, without logical reasoning being involved. Such a mechanism makes it superfluous to argue that complex logical operations cause the choice. The critical question is then: is behaviour when facing single options sufficient to predict behaviour when two or more options are present? If the answer is yes, then we should remain agnostic as to what mental operations happen at choice time. Curiously, this simple control (observing behaviour towards single options) is not reported in studies that conclude that choices are supported by the construction of preference through logical reasoning at the time of choice.

Patterns of responding to single options can have multiple origins. They may be innate or acquired through learning. These distinctions are interesting but not crucial to the present argument. What is critical is whether options faced in choices are treated in parallel rather than being evaluated through some rational process. The hypothesis of parallel processing of options has been successful in diverse protocols (15–17). The rationale is embodied in the Sequential Choice Model (SCM), that includes hypotheses about the environment of evolutionary adaptation, about individual learning and about the cognitive process of action competition in choice encounters. This is a summary:

(1) *Environment:* sequential encounters with single targets are probably more frequent and evolutionarily influential than choices between multiple options.
(2) *Acquisition:* Experience (or natural selection) leads to appropriate behaviour (pointing, grasping, pecking, touching, approach-withdrawal, sniffing, etc.) towards recognisable stimuli with predictable contingencies. For instance, a frog may innately snap at a worm and ignore a leaf, or an infant primate may learn through social feedback to gesture towards a ripe fruit and ignore an unripe one.
(3) *Choice:* if two or more targets are faced simultaneously, their option-specific actions compete for expression. The hypothetical frog will snap the worm and the infant primate will point at the ripe fruit, without constructive cognition at choice time.

These three components have different status:

The first addresses ecological relevance and is not critical to the model’s verification. It is plausible, because if potential targets have independent distributions, facing them sequentially should be more frequent and evolutionarily influential than simultaneously. Early applications of optimality theory to behavioural ecology addressed patch exploitation, diet choice and copulation duration as sequential decisions of the kind ‘stay or go away’ (18–20; see also 21,22). While leks or fruit clusters exemplify simultaneous encounters, the set of scenarios in which sequential encounters prevail is rich enough to suspect their primacy.

The second element rests on behavioural observations (e.g. 23,24,16,25–27) and should be verified in every empirical test. It is not unique to this model, but this model needs it. In laboratory tests of SCM options encountered sequentially lead to the acquisition of differential latencies to respond (faster response times to better options). This may seem intuitive and trivial, but it is in fact puzzling: under the assumption of reward rate maximisation that underlies classical foraging models there is no reason why taking any opportunity within the acceptable set should take longer than minimally possible, but models that include these latencies yield better empirical fitting than those ignoring them (17,28). Another corroboration is observed in classical (Pavlovian) conditioning. When discriminable stimuli reliably predict different outcomes, animals show differential anticipatory behaviour even if the outcomes are not contingent on the behaviours (e.g. 29,30). This matters, because response latencies or other forms of anticipatory behaviours may be sufficient to explain how animals act when facing a choice.

The third element is in fact the hypothesis under test: if choices are predictable from pre-existing behaviour, then logical reasoning is not necessary at choice time.

Thus, if behaviour towards sequential stimuli is sufficient to predict preference in a choice experiment, then the experiment does not demonstrate inferential reasoning or other high-level cognitive hypotheses such as causal understanding. Mody and Carey (13), Lauffer et al. (31) and Rescorla (32) also question the evidence for current claims for inferential reasoning, but our rationale is more radical because we argue that animals facing multiple options may not even cognise the task as a choice, but rather process options independently.

### The 2-Cups Task

The search for evidence of thinking in non-human animals can be traced to the ancient Greeks. In the 3rd century BCE, Chrysippus discussed a form of reasoning called “disjunctive syllogism” or *modus tollendo ponens*, which is summarised as “Either p or q. Not p. Therefore, q”. This deceptively simple operation assumes competence to handle the relation ‘OR’, implying that out of two world states only one can be true (see 13). It also requires understanding the concept of negation, ‘NOT’. Together, given OR, if one is negated, the other must be true. Five centuries later, Sextus Empiricus wrote that Chrysippus had used a thought experiment to claim that a dog could demonstrate logical reasoning without language, through its behaviour. In a simplified version, an imaginary dog chases a hare that gets out of sight where the road bifurcates. The dog sniffs one side and, detecting no hare smell, runs straight into the other, without sniffing. The failure to seek information in the latter is taken as evidence of inferential reasoning by exclusion. This assumes that the dog understands that there is exactly one hare, that hares do not vanish, and that the probabilities of it being in each of the roads add to unity: if it is zero in one, it must be one in the other. Following this line of thought, at a behavioural level the dog’s actions would be consistent with such logic, but reasoning is not required. If in its past whenever the dog ran into a sniffed-but-odourless road it missed dinner, but running into un-sniffed roads had mixed outcomes, it will waste no time in entering uncertain, un-sniffed, roads but ignore odourless roads. In a road fork, this would result in a ‘choice’ without the need for logical reasoning; the dog runs into the unsniffed road without using OR or NOT, does not estimate probabilities, and does not assume that there is exactly one hare in the world.

In cups tasks a subject familiar with receiving food after appropriate behaviour (such as pointing) faces two or more opaque containers (6,7,13,33). Different variations are used to investigate causal and inferential reasoning as well as more complex psychological constructs such as belief revision (see 34). Here we focus on the simplest 2-cups case, and would later argue that our arguments apply equally to most other variants (see *Value-dependent targeting of actions or Bayesian revision of beliefs?* Discussion section). The protocol starts when an experimenter visibly (to the subject) hides one bait without identifying in which cup. This is meant to inform the subject that there is one and only one bait. Then the experimenter briefly lifts one of the cups, showing its content (or absence of it), leaving the other resting. For each cup, the subject has witnessed one out of three events that signal that the cup is baited, empty, or uncertain. The subject then points to a cup and receives its content (i.e. food or nothing, see **Figure 1A**). The three possible events are thus observable stimuli that signal important outcomes. Notice that following Stevens (35) we do not restrict our notion of a stimulus to the physical energy, but to transformations of the perceived environment that cause a response. This is crucial, because, as Terrace (36) stated, “*By expanding the traditional definition of the stimulus, the study of animal learning has metamorphosed into animal cognition*”. The perception sequences that elicit a response are not two or more identical cups, but cups which have different recent history.

In the supercognitive interpretation, if the subject points at the cup that had been resting when the alternative one was exposed to be empty, it is concluded that it has cognitively performed an *Inference by Exclusion* logical operation. In the SCM interpretation, the animal would treat each of the three stimuli differently even if they were met in isolation, for instance pointing fast to a cup previously lifted and shown to be baited, slower to a resting one, and not at all to a cup lifted and shown to be empty. If the subject just aims to act in the same way when two cups are present, it would display the correct ‘choice’ without logical reasoning.

Preference for the baited cup in *2-Cups* protocols can be generated by multiple processes. M. Rescorla (32) used a Bayesian analysis to argue that *“results supposedly indicative of non-linguistic deduction can be explained without citing logical reasoning or logically structured mental states”*, and Lauffer and colleagues (31) used a 2-alternative forced-choice discrimination task to demonstrate that behaviour consistent with inference by exclusion, could also be explained by acquired equivalence, an associative rather than reasoning process. Both are algorithmically efficient explanations that omit ‘logical reasoning’ or ‘causal understanding’. We go further, and demonstrate experimentally that animals do solve such tasks without even framing the problem as a choice.

### An emulation of the 2-Cups Task

Our experimental procedure reproduces the critical contingencies between stimuli and behavioural outcomes in the *2-Cups Task* (**Figure 1B**). Starlings went through three phases: pre-training, sequential training, and preference test. They were first pre-trained to peck at four different stimuli encountered sequentially as single options. In the *Sequential Training* phase three stimuli were paired to different contingencies, labelled *Sure Reward* (emulating an exposed baited cup), *No-Reward* (emulating an exposed empty cup), or *Uncertain* (emulating a cup left resting). In each trial, a response to a central attention key was followed by a Pre-Go period in which one of the three stimuli appeared for 2s on either of the two side keys. During these 2s, pecks were recorded (as ‘Pre-GO Pecks’) but had no programmed consequence. Once the 2s lapsed, the stimulus was switched off and a ‘GO’ stimulus (always white) was presented in the same side key. The first peck at the GO key realised the trial’s food contingency (**Figure 2A**, top). After 10 days in this phase, the *Preference Testing* phase followed. This phase included choice trials with both side keys showing two of the three stimuli simultaneously for 2s and then turning white to enter the GO state. A peck to either key in the GO state caused the offset of the alternative and the realisation of the respective food contingency (**Figure 2B**, top). Pre-GO pecks were recorded but had no effect. They served to test whether behaviour in choice trials reproduced that in sequential ones. Choices were determined by the first peck to a key in the GO state. This emulates choosing between cups by pointing at one in the *2-Cups* protocol. We describe starlings’ behaviour in sequential encounters and in choices, examining whether responding in single encounters is sufficient to predict choices in simultaneous trials.

**Figure 2.**
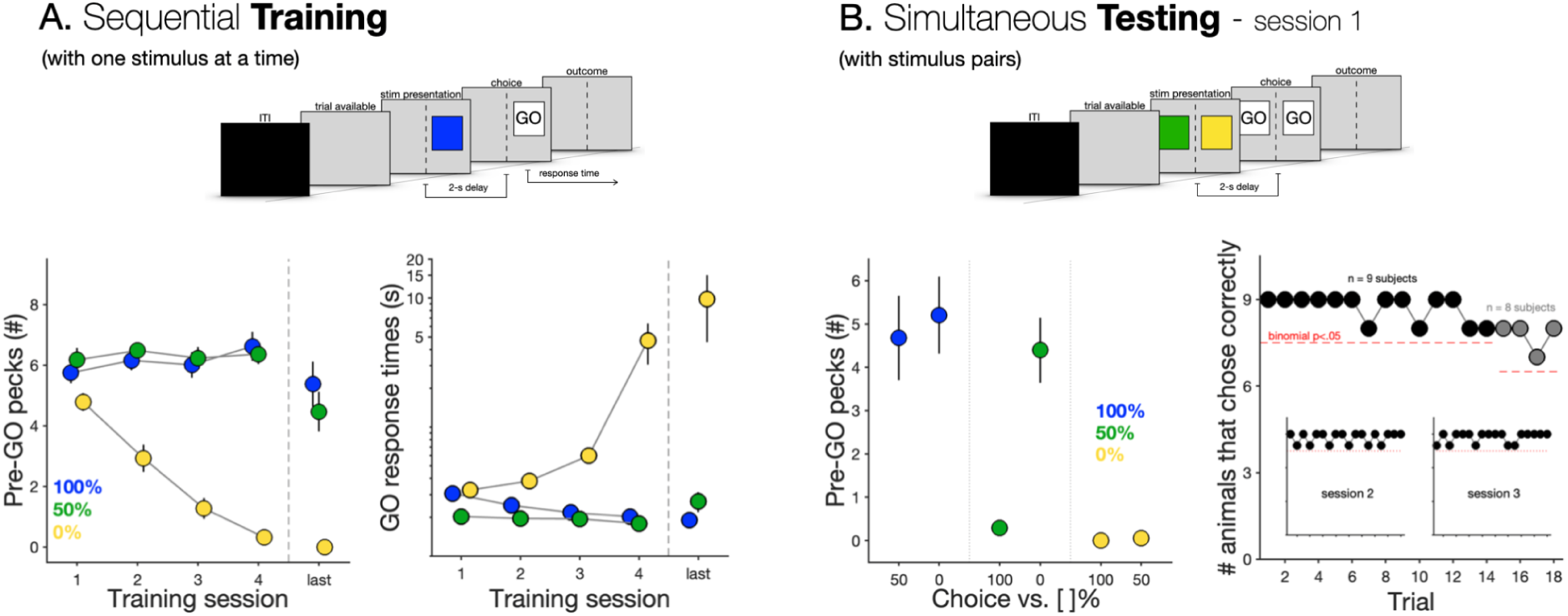
Starlings’ performance in the 2-cups task analogue. (**A**) Example of a sequential trial (top). Average (±SE) Pre-GO pecks (bottom left) and response times (bottom right) for each alternative (*Sure Reward* - 100%; *No Reward* - 0% and *Uncertain* - 50%) during the first 4 (and last) sessions of sequential training. (**B**) Example of a simultaneous choice trial (top). Average (±SE) Pre-GO pecks for choice trials for all simultaneous choice trial types: *Sure Reward* vs. *Uncertain*, *Sure Reward* vs. *No Reward*, *Uncertain* vs. *Sure Reward*, *Uncertain* vs. *No Reward*, *No Reward* vs. *Sure Reward* and *No Reward* vs. *Uncertain*, respectively in blue, green and yellow (bottom left). Number of animals choosing *Uncertain* (trial-by-trial) in choices between it and *No Reward* for the first testing session (see Figure S5 for the remaining choice pairings). Black and grey markers depict trials with 9 and 8 subjects, respectively, and red dashed line binomial significance. Insets show the same data for testing sessions 2 and 3 (n=9 subjects).

Our procedure emulates the contingencies in the basic *2-Cups* protocol, with arbitrary stimuli and controlled acquisition instead of food containers. Subjects in *2-Cups* experiments do not have a controlled pre-training phase, but they have enough experimental experience to ‘know’ that a container recently seen to be full is likely to contain food and one whose inside remains unseen can be either full or empty, hence are likely to respond differently when facing each scene. This difference in training procedure between the *2-Cups* experiments and our emulation is not an obstacle for the present argument. What is critical is whether behaviour in sequential encounters is enough to predict preferences, regardless of how differential responding is acquired. If reports of *2-Cups* experiments included the behaviour of subjects in single cup presentations, the choice-without-reasoning interpretation would be either confirmed or rejected.

## Results

The hypothesis under test is that choice (pecking at one of the GO stimuli in a choice trial, equivalent to primates pointing after the demonstration or different choice actions in other species) is pre-determined before pairs of options are encountered, implying that preference does not depend on cognitive operations that include the logical disjunction ‘OR’. We first examine behaviour during the *Sequential Training* and then in the *Simultaneous Testing* phases (**Figure 2**).

### Sequential Training phase

At the start of sequential training all starlings reliably pecked at all keys. Figure 2A shows averages of pecking at the Pre-GO stimuli during the 2s Pre-GO interval and of response latencies towards the GO key in the first four sequential sessions and in the last one, just before switching to the *Simultaneous Testing* phase. The birds rapidly extinguished responding at the *No Reward* option but maintained responding when there was a chance of food. These trends were statistically reliable. The three options were affected differently by learning. For the *Sure Reward* option, Pre-GO pecks did not change significantly (effect of session: p = 0.114), and response latency to GO decreased (p < 0.001). Relative to the *Sure Reward option*, Pre-Go pecks decreased and response times increased for both the *Uncertain* (p = 0.031 and p < 0.001 respectively) and *No Reward* (p < 0.001 for both) options. See *Supplementary text* and Table S1 — Models 1.1 and 2.1— for statistical details. Consistently, response times to the first peck at the GO stimulus remained very short for the two rewarded stimuli and increased for the unrewarded one.

By the last session of the *Sequential Training* phase both Pre-GO pecking and response times to peck at the GO key were significantly different between the *Sure Reward* stimulus and the *No Reward* one (p<0.001 for both pecks and response times), with no significant differences between the two rewarded alternatives (pecks: p=0.086, response times: p=0.14; see Table S1 — Models 3 and 4— for details and figs. S1 and S2 for individual results). In terms of the *2-Cups Task* emulation, that would be equivalent to subjects discriminating behaviourally between the unrevealed uncertain cup and the revealed empty one (in this emulation the *Uncertain* and *No Reward* arbitrary stimuli) when a single cup is presented.

### Simultaneous Testing phase

In the final, preference phase, subjects faced a pseudo-random sequence of ‘sequential’ and ‘choice’ trials (see *Materials and Methods*, “*Simultaneous Testing*” phase subsection). We focus on preference during Pre-GO between *No Reward* and *Uncertain*, as this emulates the critical choice between a lifted empty cup and a resting cup, the condition used as evidence of inference by exclusion.

### Pre-GO pecks in choice trials

Figure 2B (bottom left; see also Figure S3 for individual data) shows Pre-GO pecks in the first choice session, in three panels. The left panel shows Pre-GO pecks at the *Sure Reward* (100%) stimulus when paired with either the *Uncertain* (50%, left) or *No Reward* (0%, right) alternatives. In both cases pecking did not differ from that in sequential trials (p=0.766 and p=0.064, respectively; see Table S1 —Models 6 and 7—for details). The middle panel shows Pre-GO pecks at the *Uncertain* (50%) stimulus when paired with the *Sure Reward* (100%, left) or *No Reward* (0%, right) alternatives. In this case, Pre-GO pecks depended on the alternative. When paired with *Sure Reward* (100%) pecking dropped significantly (p<0.001; see Table S1 —Models 8— for details), while when paired with *No Reward* (0%) pecking was maintained (p=0.239; see Table S1 —Models 9— for details). This is evidence for response competition: the animal cannot be in two places at once and discrimination ability acquired in sequential encounters is not fully captured by the number of pecks in sequential encounters, probably due to a ceiling effect. Finally, the rightmost panel shows Pre-GO pecks at the *No Reward* (0%) option when paired with either of the other two. In both cases there was very little pecking.

In summary, Pre-GO pecking, although inconsequential in terms of food delivery, demonstrates that discriminating behaviour exists before the animal is allowed to choose, in the sequential phase (for Pre-GO pecking in sequential trials during the Simultaneous Testing phase see Figure S4). In terms of the *2-Cups Task* emulation, a direct comparison between the experiments could be explored if subjects’ behaviours such as gazing or pointing were reported during the demonstration or just before consequential (instrumental) choices.

### Choice in paired trials

Successful subjects in the *2-Cups Task* prefer the resting cup when paired with a lifted but empty one. Figure 2B (bottom right) shows that also starlings were almost totally error-free in this test from the very first choice trial (this persisted throughout this session and the following ones). The mean proportion of correct choices in the first choice session was 0.968, ranging from 0.951 to 0.980 when subtracting and adding the standard error, respectively, this being substantially better than random (p<0.001; see Table S1 —Model 5— for details). The birds could have achieved this without logical inferences, by treating each stimulus as a competing opportunity to respond. The remaining pairings, although less relevant to the arguments, are presented in detail in the *Supplementary Materials*.

## Discussion

### Simple cognitive processes can lead to rich and adaptive behaviour

Because simple cognitive processes can lead to rich and adaptive behaviour, the presence of such behaviour is not evidence of complex cognition. We illustrate this argument by showing that differential actions towards behavioural targets met in sequential encounters generate correct choices when multiple opportunities are met simultaneously. We believe this mechanism to be very widespread and very important for the interpretation of results in choice experiments. The specific mechanism we explore with starlings shows that they behave consistently with reasoning by exclusion, but their choices are a consequence of each stimulus in simultaneous sets eliciting behaviour acquired in sequential encounters. Thus, behaving as if making inferences by exclusion does not require forming estimates of probabilities or connecting alternatives with the OR disjunction and can result from processes less computationally demanding than logical reasoning.

In addition, this mechanism seems ecologically more relevant, since encounter rates with single potential targets in nature are probably more frequent than simultaneous encounters requiring a choice. It is possible that many species have evolved mechanisms to behave appropriately and differently to different stimuli but have no dedicated mechanisms for choosing between simultaneously present targets.

We provide a novel test of the Sequential Choice Model, which in essence states that behaviour in sequential encounters can explain choices in a variety of experimental protocols (15–17). Although some may consider this to be a ‘killjoy’ approach (10) given the widespread appeal of attributing high-level cognition, the opposite view is compelling: simple mechanisms of behaviour can be readily implemented in natural or artificial agents and are sufficient to produce rich, flexible, and adaptive behavioural repertoires. When sufficient to explain the observations, they should be preferred for their parsimony and simplicity. This is encouraging not only because it advances understanding of animal behaviour, but also because it potentially simplifies the design of intelligent artificial agents.

There are important caveats regarding the extent to which our experiment constitutes an emulation of the *2-Cups* protocol. We explicitly trained the starlings with sequential encounters before giving them choices, while this is not the case in tests of the *2-Cups* task. We believe that this is not a fatal difference for two main lines of argument. First, all animals tested in *2-Cups* protocols have rich histories, in which they can have acquired the relevant competences. In a study interpreted as a demonstration of causal understanding in chimpanzees, Hanus and Call (2) wrote:

*“Recurrent experience during ontogeny definitely plays an important role in the acquisition of such cognitive skills—in human and nonhuman primates alike. Individuals need to interact repeatedly with various causal regularities within their physical world in order to be capable of transferring this specific knowledge to new situations.”*

This caveat applies to *2-Cups* experiments because individuals must learn to treat a cup lifted and shown to hide a bait differently from one that is resting on a surface. If a subject solves the *2-Cups* problem as if using inferential reasoning, it will most likely respond differently to the same cups if it were tested in single-option sequential presentations. This is all that is needed for our emulation to be informative. We focused our analysis on the very first choice faced by the starlings, to exclude the possibility that their behaviour had been shaped by having been rewarded for particular choices. On their first simultaneous presentations the birds had no other way of choosing correctly (as they did) than relying on biases acquired in sequential encounters, when no inferential reasoning by exclusion is relevant. Our second, most conclusive argument is that lingering doubts about the relevance of our emulation have a clearcut empirical solution by adding a control to the *2-Cups* protocol: testing what subjects do when a single cup, either shown to be baited/empty or not exposed, is presented, namely by being briefly lifted or left resting. If, as we expect, their behaviour towards single targets predicts what they do when facing two cups, then there would be no need to invoke a constructive cognitive operation such as logical inference at the time of choice. We look forward to reports of such controls.

### Value-dependent targeting of actions or Bayesian revision of beliefs?

We discussed and emulated the simplest version of the cups protocol, but in recent years additional controls and more complex designs have been implemented (e.g. 13,33). For instance, Schleihauf and colleagues (34) reported multiple experiments that led them to argue that “*chimpanzees metacognitively evaluate conflicting pieces of evidence within a reflective process”.* Their experiments showed that chimpanzees facing two consecutive choices did not rigidly respond according to the first paired display but instead modified their behaviour as a function of what they saw in both demonstrations. Their article was titled “*Chimpanzees rationally revise their beliefs*”, and was later commented under the title “*Chimps can weigh evidence and update their beliefs like humans do*” (45; see also 46). However, the reported results could have been achieved through the process we discussed above. What is necessary is that chimps attribute value (utility) to each event-stimuli and express different behaviour such as faster uptake (38) when targeting each class of event-stimulus (Table S2 describes how this would work for all of Schleihauf et al.’s experiments). As in the simpler *2-Cups* protocol, our argument can be explored by reporting behaviour towards each stimulus when presented alone. If they did show differences such as adjustments of reaction times towards single stimuli, parallel expression of such behaviour would suffice to predict all results.

We believe that it is not necessary to fall in the restrictive framework of radical behaviourism to recognize the weaknesses of supercognitivism, which overlooks the virtues of parsimony. The obtained results seem impossible to explain without assuming value representation, an element that underlies choice transitivity, so employing such constructs seems justified. However, higher-level inferred attributes, such as hypotheses or belief modification, inferential reasoning or causal understanding do not seem unavoidable given present evidence and should therefore be invoked with restraint.

A different argument that is often raised can be summarised as “*if humans do it, why wouldn’t other species?*”. Leaving aside that much of human choice mechanism is likely not to be based on logical reasoning, this line of reasoning makes the hypotheses of other species’ possession of sentience, consciousness, and logical reasoning capabilities entirely plausible, but it does not justify the claim that such capacities have been demonstrated by the available evidence.

## Materials and Methods

### Subjects

Wild-caught adult spotless starlings (*Sturnus unicolor*; n=9) were housed in pairs in indoor cages (1350 × 784 × 800 mm (l × w × h)) and kept under a 12L:12D cycle with gradual transitions at dawn and dusk. They were confined individually to one section of their cage for tests, but after each session, they had 4h of ad-libitum food and social interaction with the cage-partner. By the end of the testing season (early summer), the birds were released back into the wild.

The experimental protocols were approved by the University of Aveiro Animal Welfare Body (B003/2020) and by the Portuguese General Directorate of Food and Veterinary Medicine (0421/000/000/ 2021), and their capture and holding were approved by licenses 514/2021/CAPT and 07/2021/DET respectively (both from the Portuguese Department of Wildlife and Conservation).

### Apparatus

The experimental cages had two side areas that could be split with sliding panels from a shared middle section (see 37). Each side area had a working panel with three sections, each with a centrally placed response key (**Figure 1B**). The central section had a food hopper connected to a pellet dispenser (Campden Instruments) containing 20 mg *BioServ* precision dustless pellets. Contingencies were controlled by custom software running on the Microsoft WINDOWS operating system and attached to an Animal Behaviour Environment Test System (Campden Instruments) via WhiskerServer.

### Trials and Experimental phases

The experiment had two phases: S*equential Training* and *Simultaneous* (pairwise) *Testing*. During the *Sequential Training* phase only single-option trials took place, while in the *Simultaneous Testing* phase both sequential (single-option) and choice (two options) trials were interspersed. In sequential trials, a pre-GO stimulus was presented for 2s, followed in the same location by a GO stimulus, always white, that stayed on until pecked once (**Figure 2A**, top). In choice trials two pre-GO stimuli were simultaneously presented (**Figure 2B**, top), each followed by a GO stimulus. The stimuli were square 4 by 4 LED light panels lit in green, blue, yellow or white (**Figure 1B**). The three colours were used as pre-GO stimuli, and white served as a common GO stimuli that, when pecked, realised the contingency. The three discriminative contingencies were *Sure Reward* (food delivered with 100% probability), *No Reward* (food delivered with 0% probability), and *Uncertain* (food delivered with 50% probability). During the *Simultaneous Testing* phase pre-GO stimuli were paired in their three possible combinations plus a fourth double presentation of the *Uncertain* stimulus in both left and right key (see *Simultaneous Testing* phase for details).

### General procedure

Each trial began with the centre key displaying a blinking (700 milliseconds ON, 300 milliseconds OFF) white square, signalling that a response was available. Blinking continued until the subject pecked the centre key, which turned it off and was followed by either one (sequential trial) or both (simultaneous choice trial) side keys being lit and the 2s Pre-GO period starting. In sequential trials (**Figure 2A**, top), the unlit key remained disabled. In choice trials (**Figure 2B**, top), both side keys lit up and were enabled. Depending on the trial type, the stimulus was either an unblinking green square, or a blinking (700 milliseconds ON, 300 milliseconds OFF) blue or yellow square (see the “*Sequential Training* phase” sub-section). During these stimuli (2 seconds; referred to as the “Pre-GO period” hereafter), pecks to side keys had no programmed effect but were recorded and later served as dependent variables.

At the end of the Pre-GO period, illuminated side key/keys (one or two in sequential or choice trials, respectively) switched to white (the GO signal) and remained lit until pecked. Food was delivered or not depending on the corresponding Pre-Go stimulus. After this, both keys were turned off again for a 30-s inter-trial-interval (ITI).

Trials terminated if the subject failed to peck at the GO key within 1 minute. If the stimulus had been *Sure Reward* or *Uncertain*, after an ITI the same trial was repeated, up to a maximum of 3 presentations (correction procedure).

### Pre-training

All subjects had experience with the apparatus in tasks with different contingencies (studies of ‘suboptimal/paradoxical choice’, and ‘mid-session reversal’). In the *Pre-training* phase, pecking at the white central key was followed by the onset in a side key of one of the stimuli that would later serve in the sequential and choice phases. All keys were programmed with a fixed ratio (FR) reward schedule. Over two days, each stimulus was presented for three 50-trial sessions, divided in blocks of 25 trials with a 15-minute break and with a 60-s ITI. The fixed ratio increased in consecutive sessions, with FR1, FR3 and FR5 in the first, second and third sessions respectively. One individual needed an extra session due to apparatus malfunction. Before starting the Sequential Training phase, subjects had an additional FR5 session consisting of two blocks of 35 trials for each of the 4 stimuli. At the end of pre-training, all subjects were pecking reliably and similarly to the three coloured and white stimuli.

### Sequential Training *phase*

In the Sequential Training phase, only one alternative was presented in each trial, while the other key remained dark. The side of the stimulus presented was pseudo-randomized so that it appeared the same number of times on both sides and no more than twice consecutively on the same side.

One of three coloured stimuli was presented in each trial, each associated with a specific probability of obtaining a food reward (Figure 1B; see also 38). For half of the subjects the Sure Reward stimulus was yellow and the No Reward stimulus was blue, and the opposite for the other half of the animals. For the *Uncertain* stimulus, the key stayed green for the 2 seconds of Pre-GO time.

The subjects were trained daily, with up to 4 sessions per day, each comprising 108 trials. For 5 animals, the first training day comprised 3 sessions (at that point it was unknown whether the animals would complete 4 sessions within a day). Hence, analyses of their first 4 sessions include the first session of their second training day.

All subjects shifted to the *Simultaneous Testing* phase after 10 consecutive days in sequential training.

### Simultaneous Testing *phase*

During Simultaneous Testing, sequential trials were interspersed with five kinds of simultaneous choice trials, in which two alternatives were presented, one in each side key.

There were 4 different choice trial types that emulated the possible contingencies in a 2-Cups procedure, as follows:

1. *<u>Uncertain</u>*vs. *<u>No Reward</u>*: This choice emulates trials in the *2-Cups* procedure in which one cup is left resting and the other is lifted briefly to show that it is empty. Correct choice with these contingencies is interpreted as showing Inference by Exclusion.
2. *<u>Sure Reward</u>* vs. *<u>No Reward</u>*: This emulates a choice between a cup shown to be baited and another shown to be empty. This trial type came in two variants with stimuli either presented simultaneously (synchronous) or sequentially (asynchronous), to control for recency effects. In asynchronous choice trials the second stimulus started after the end of the 2-second Pre-Go time of the first one. The apparatus remained unresponsive until the end of the second pre-Go period. In the case of the *asynchronous* variant, the order and side of stimuli were balanced.
3. *<u>Sure Reward</u>* vs. *<u>Uncertain</u>*: This emulates a choice between a cup shown to be baited and another left resting so that its contents are unknown by the subject. In these trials Uncertain were never programmed to deliver food if chosen. Because our analysis is mainly based on preference in the initial trials of the choice phase, when this subcontingency had not yet been experienced and (as described under Results) choice was virtually exclusively towards *Sure Reward*, it had no opportunity to exert any influence.
4. *<u>Uncertain</u>* vs. *<u>Uncertain</u>*: The purpose of these trials was to control for side preferences when both contingencies are equated.

Trial types were presented in a pseudo-randomized manner, with no more than two consecutive trials of the same kind or where the expected correct response was on the same side. Sessions in this phase had 90 choice trials (18 per pairwise combination, with *Sure Reward* vs. *No Reward synchronous* and *asynchronous* counting as two separate trial types) and 18 sequential trials (6 per pairwise combination). This phase lasted for 10 consecutive days. An experimental error in Uncertain vs. Uncertain trial type during the first testing session of the first 5 animals being tested led to an early abortion of this session for the corresponding animals (leading to the early dropout in subjects seen in the last few trials of Figures 2 and S5). Due to this error, for the 5 animals affected, the data used for analyses and graphs not involving the Uncertain vs. Uncertain trial type starts at session 1, while for the graph involving this trial type (last panel of figure S5), data from session 2 was used instead. All corrupted data was removed from the dataset provided. Pseudo-randomization rules applied equally to sequential and choice trials.

### Statistics

We analysed the number of pecks in the Pre-GO periods and, concerning the GO stimuli, both proportion of choices and time to the first peck, which resulted in a ‘choice’. The main topic of interest is how behaviour in the *Sequential Training* phase relates to behaviour in the initial trials with simultaneous choices, when rules for logical reasoning about interdependencies such as ‘if the reward is not here it must be there’ are implausible because animals had never yet experienced simultaneous contingencies.

#### Acquisition of discrimination

To examine acquisition of responding as a function of experience, response times to the GO stimulus during the first day of the *Sequential Training* phase were analysed through a Linear Mixed Model (LMM) with the logarithm of the response times as the response variable, the subject as a random intercept, and the identity of the stimulus (*Sure Reward*, *No Reward*, or *Uncertain* stimuli) as both a fixed effect and a random slope. To calculate any changes in response times during the 4 sessions of the first day, one model (see **Table S1 — Model 1.1**) included the interaction between the stimulus and the session number (z-transformed, added as a random slope as well) and another without this variable (see **Table S1 — Model 1.2**). The same model structure as this latter one was used to analyze any differences in response times between stimuli in the last day of the *Sequential Training* Phase (see **Table S1 — Model 3**).

Since the amount of interactions with the keys before the GO stimulus was also dependent on the animal’s acquisition of the stimuli (with more pecks for preferred stimuli), a parallel set of Generalized Mixed Models (GLMM) with a Poisson distribution were run with the same model structure but with the number of Pre-GO pecks as a response variable (see **Table S1 — Models 2.1, 2.2, and 4**). Due to the distribution of the data, both models accounted for zero-inflation.

#### Choices in simultaneous trials

In order to ascertain whether the animals significantly preferred the rewarded option in the *Uncertain* vs. *No Reward* trials (a behaviour that would be consistent with *Inference by Exclusion*), the outcome (success or failure) of the trials was analysed using a GLMM with a binomial distribution and a logit link function, with subject as a random intercept. As the purpose of this model was to calculate the difference between the observed results and chance (a probability of 0.5), no fixed effects were added to the model, and calculations were made based on the intercept (as per 39; See Table 1 — Model 5).

To assess any potential changes in the amount of Pre-GO pecks for each stimulus between the *Sequential Training* and the *Simultaneous testing* phases (and thus, any changes in the animals’ response as soon as more than one alternative was available), an additional set of GLMMs with a Poisson distribution was run using data from the last session of the *Sequential Training* phase and the choice trials of the first session of *Simultaneous Testing* phase. A different model was fit for the stimuli in each of the potential choices (*Sure Reward* vs. *Uncertain*, *Sure Reward* vs. *No Reward*, *Uncertain* vs. *Sure Reward*, and *Uncertain* vs. *No Reward*). All of these models had the phase (*Sequential Training* or *Simultaneous Testing*) as both a fixed effect and a random slope, and the identity of the subjects as the random intercept (see **Table S1 — Models 6 to 9**). Due to general lack of Pre-GO pecks for the *No Reward* stimulus, no models were fitted for this.

All statistical analyses were carried out in R with the packages lme4, lmerTest, and car (40–44).

## Acknowledgments

We would like to thank Jeff Stevens, Sandro Sehner, Christian Menne and Robert Hampton for fruitful discussions on earlier versions of the manuscript, and Ed Wasserman, Leyre Castro, Ralph Miller and Tom Zentall for feedback on the current version. **Funding:** D.R.B., F.R. and T.M. were supported by the Austrian Science Fund (FWF) Grant DOI 10.55776/P37052 and 10.55776/P33928. D.R.B. was funded by a Marietta Blau Grant from the Austrian Agency for International Cooperation in Education, Science and Research (MMC-2023-07030). MV was supported by grant 2022.04861.PTDC from the Portuguese Foundation for Science and Technology (FCT). AK is grateful for the support of the Deutsche Forschungsgemeinschaft (DFG, German Research Foundation) under Germany’s Excellence Strategy – EXC 2002/1 “Science of Intelligence” – project number 390523135. FCT (https://www.fct.pt/en/) supported this work, through a grant to T.M. (CDL-CTTRI-249-SGRH/2022), and multiannual funding to the WJCR in the context of the R&D Unit UID/04810/2025. The funders did not have a role in the study design, data collection and analysis, decision to publish, or preparation of the manuscript.

## Author contributions

Conceptualization: DRB, FR, AK, TM. Methodology: DRB, TM. Investigation: DRB, TM. Visualization: DRB, TM. Funding acquisition: DRB, MV. Project administration: DRB, TM. Supervision: MV, FR, TM. Writing – original draft: DRB, AK, TM. Writing – review & editing: DRB, MV, FR, AK, TM.

## Competing interests

Authors declare that they have no competing interests.

## Data and materials availability

All data and code will be made available upon acceptance or at reasonable request to the corresponding authors.

## Supplementary Materials

### Supplementary text

For the sequential training, the *Uncertain Reward* Pre-GO pecks decreased (p = 0.031) and response time increased (p < 0.001) when compared with the *Sure Reward* option (the model’s intercept). However, no differences between stimuli were found at the last sequential training session (0.09 and 0.14 respectively). In order to make the informative options (*Sure Reward* and *No Reward*) more salient to the animals, we made it so that these two stimuli were blinking, while the *Uncertain Reward* stimulus was not. However, this made it so that, overall, the response key was always on for longer for the *Uncertain Reward* than for both the *Sure Reward* and *No Reward*. Given that the starlings had previous experience with the panels being active only when lit up, this may have caused them to press them less frequently (and therefore, have a lower response time on average as well) during the first few trials.

### Supplementary figures and tables

**Fig. S1.**
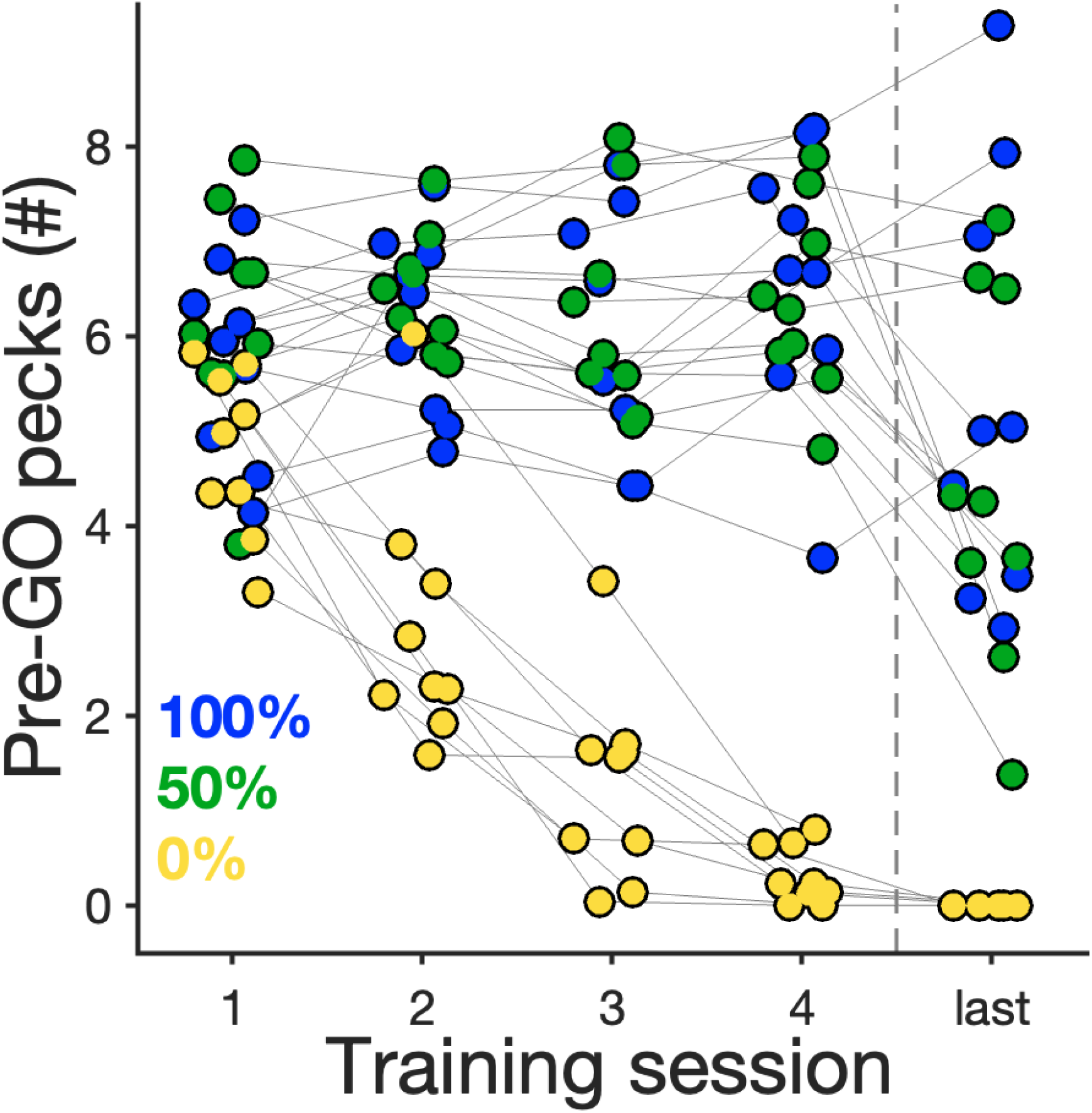
Individual average Pre-GO pecks during the first 4 (and last) sessions of *Sequential Training*. Same data as presented in Figure 2A (bottom left).

**Fig. S2.**
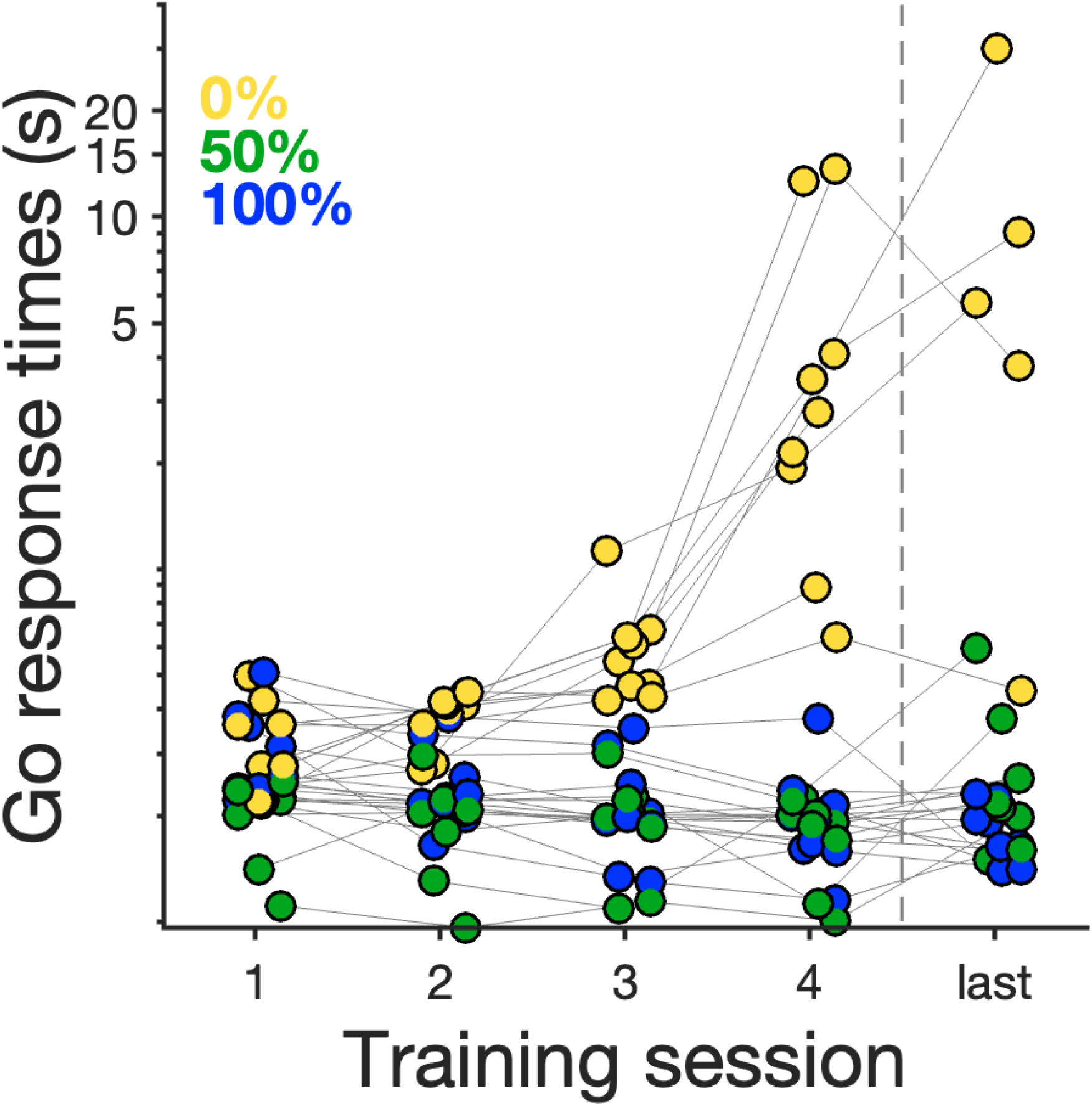
Individual median response times to the Go cue during the first 4 (and last) sessions of *Sequential Training*. Same data as presented in Figure 2A (bottom right).

**Fig. S3.**
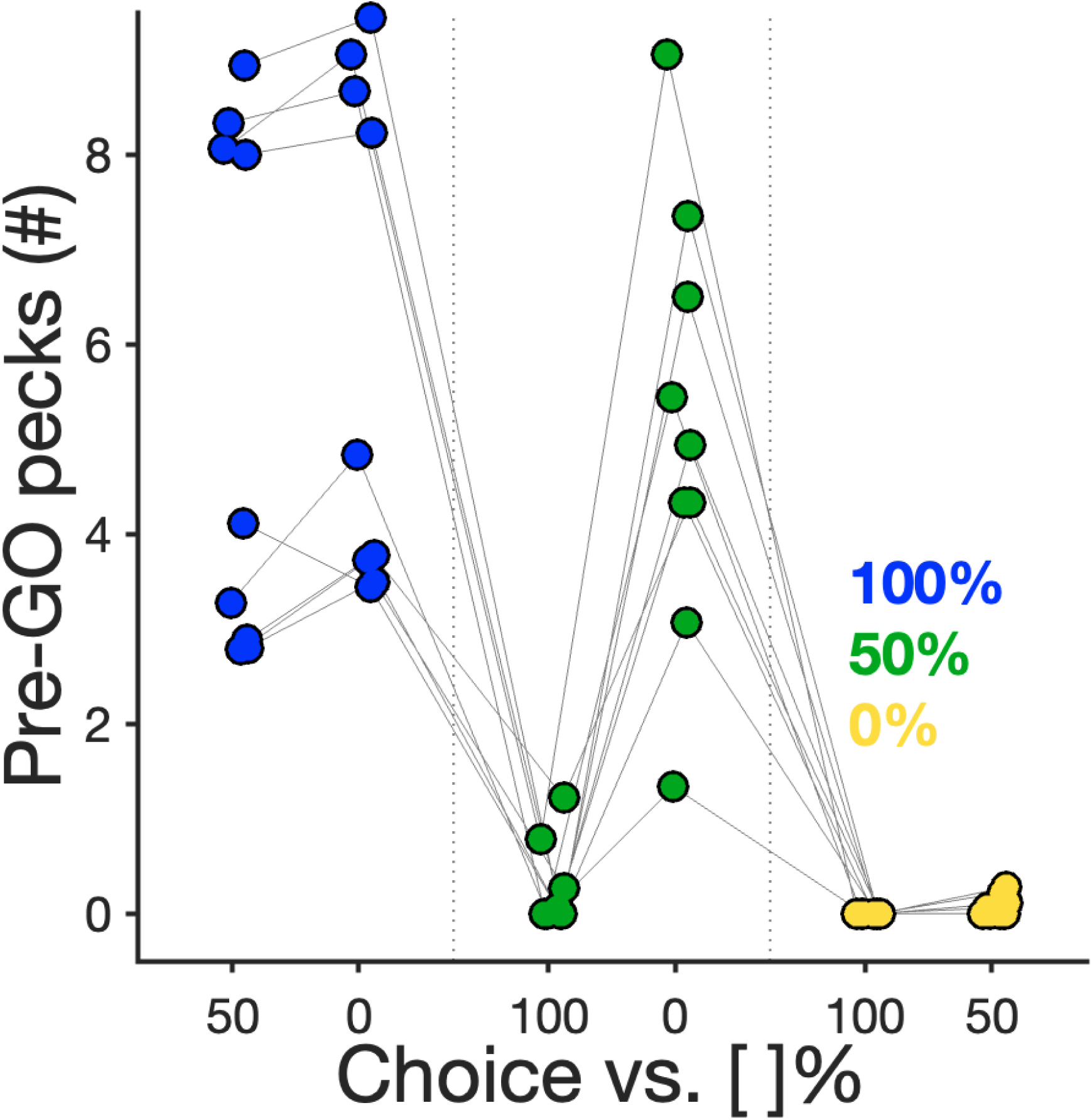
Individual average Pre-GO pecks during the first session of *Simultaneous Testing*. Same data as presented in Figure 2B (bottom left).

**Fig. S4.**
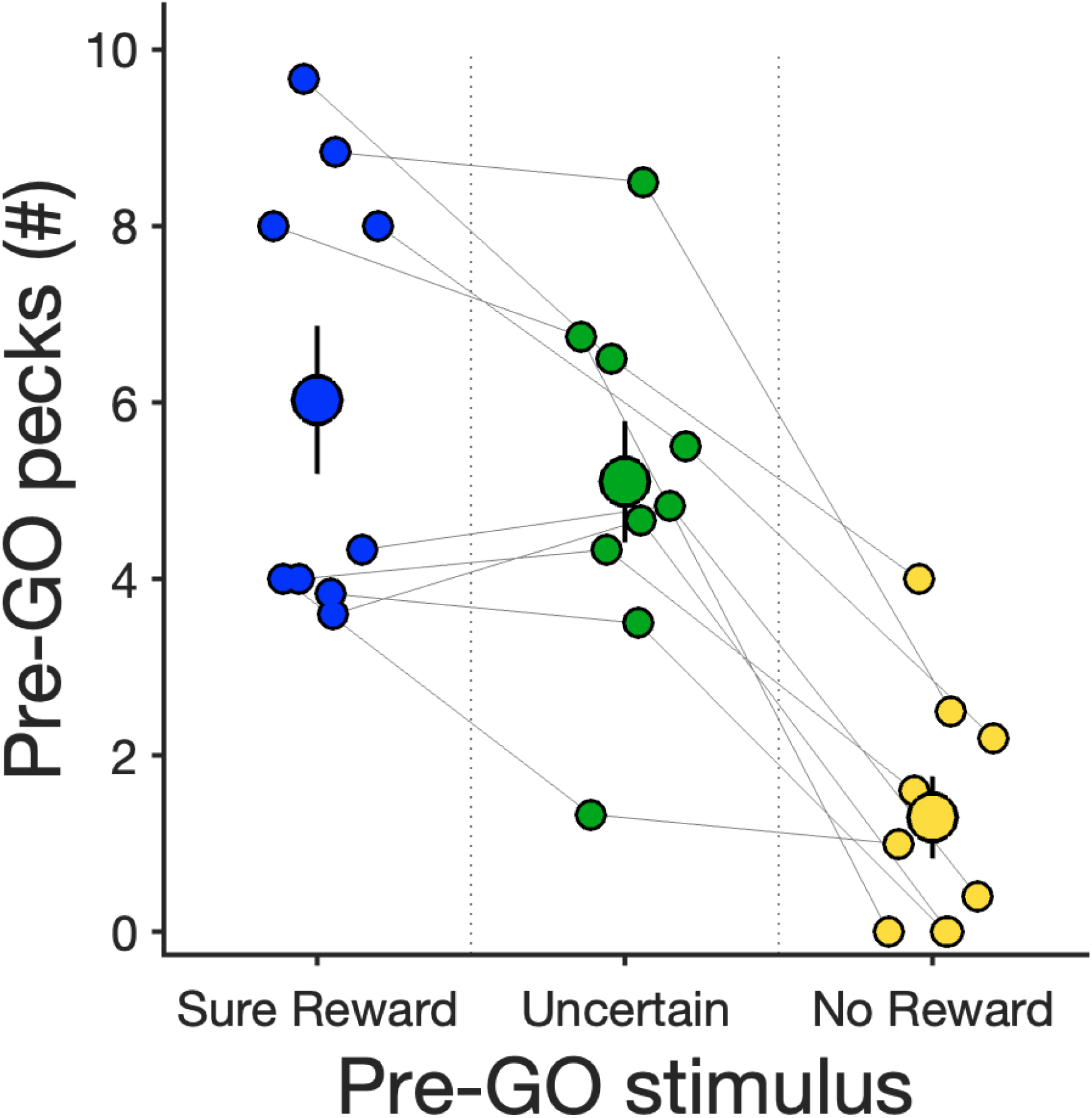
Pre-GO pecks (mean±SE) for sequential trials during the first session of *Simultaneous Testing*. Smaller markers depict individual averages.

**Fig. S5.**
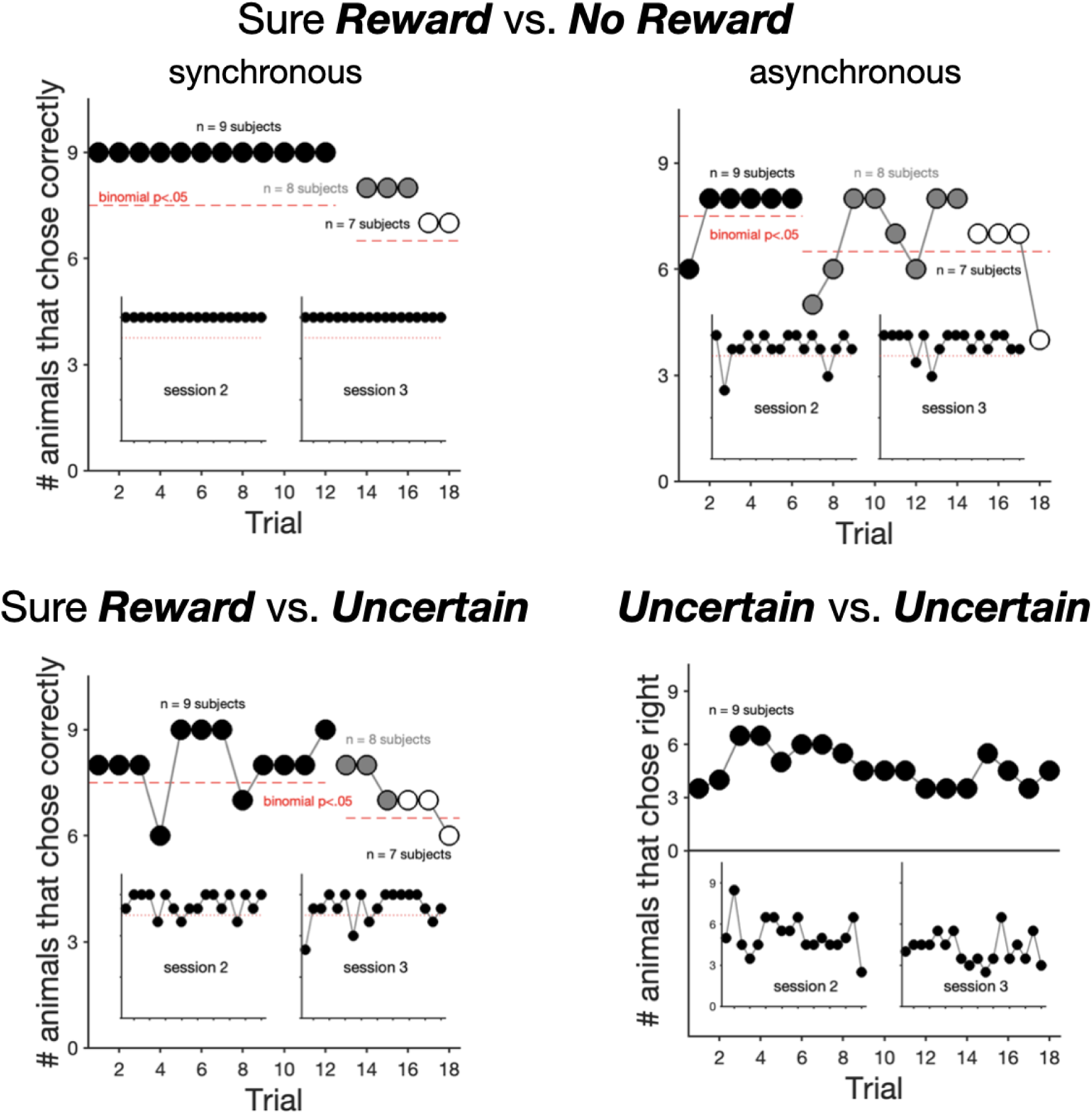
Trial-by-trial number of animals responding correctly for the first session of *Simultaneous Testing* across the simultaneous choice trial types not shown in Figure 2B (bottom right). Marker colour depicts the number of subjects tested (black=9, grey=8 and white=7 subjects, respectively. Insets show the same data for testing sessions 2 and 3. Red dashed line depicts binomial significance. For *Uncertain* vs. *Uncertain* choice type, the y-axis represents the trial-by-trial number of animals that chose the right key, as there was no single correct (rewarded) response.

**Table S1.**
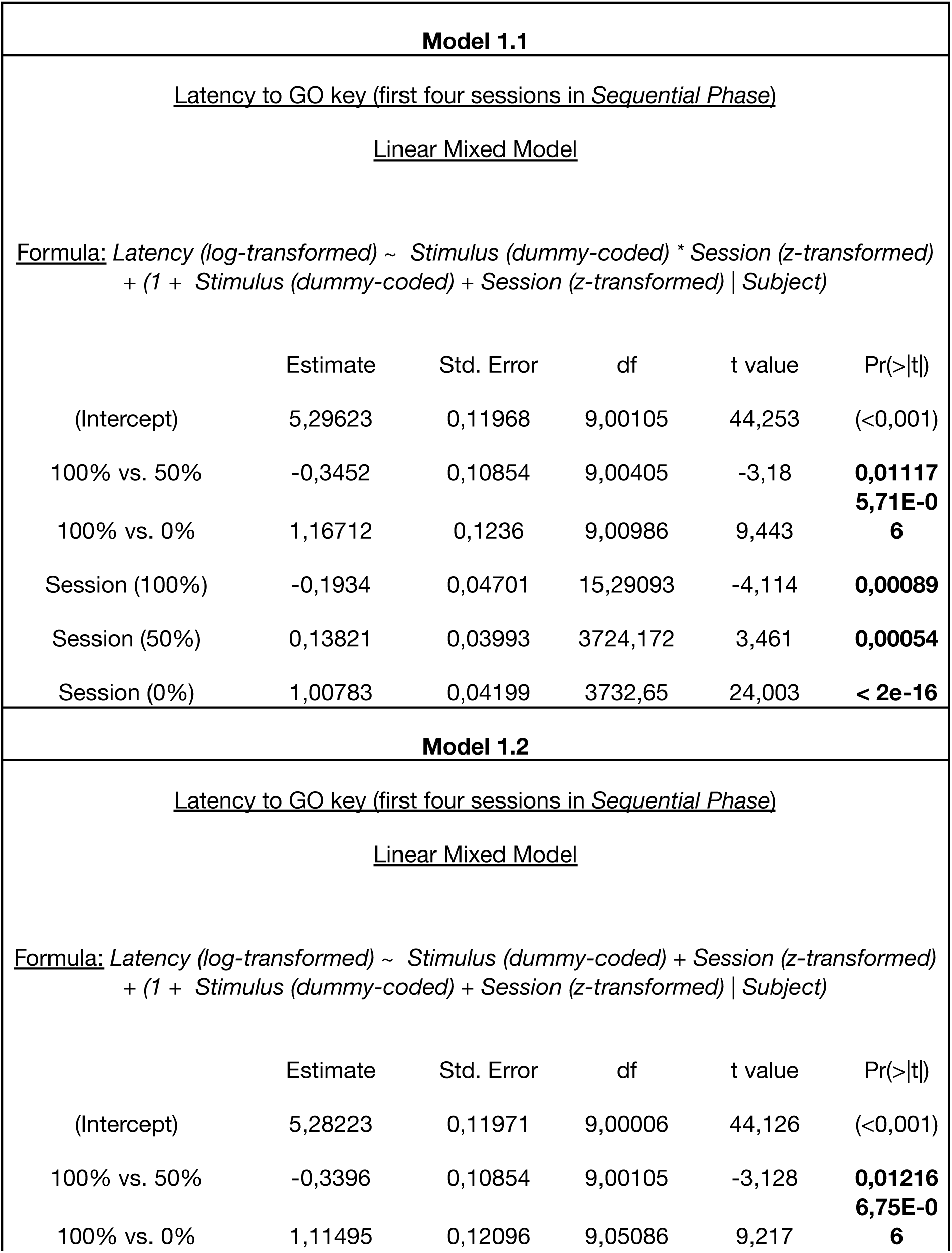
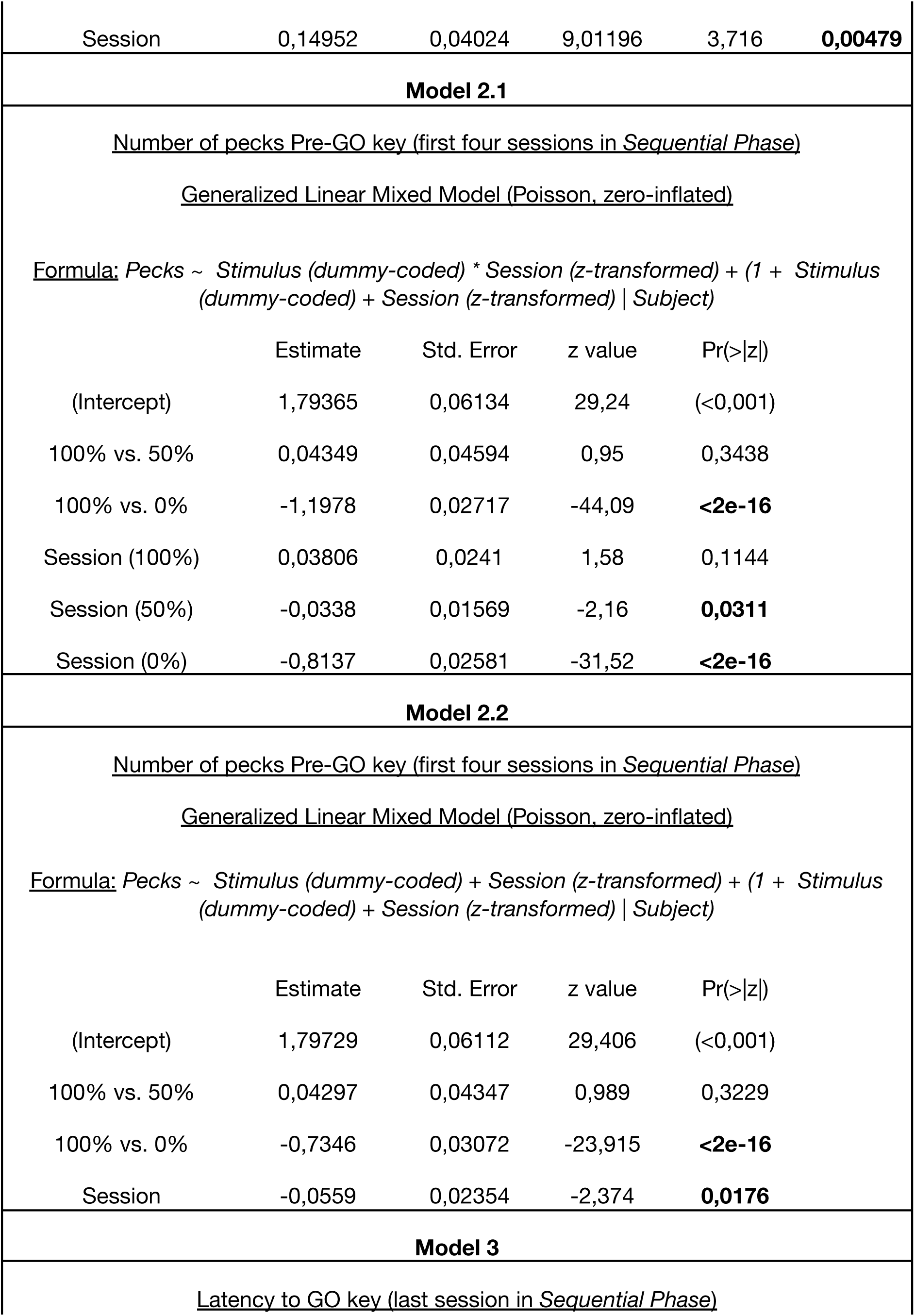
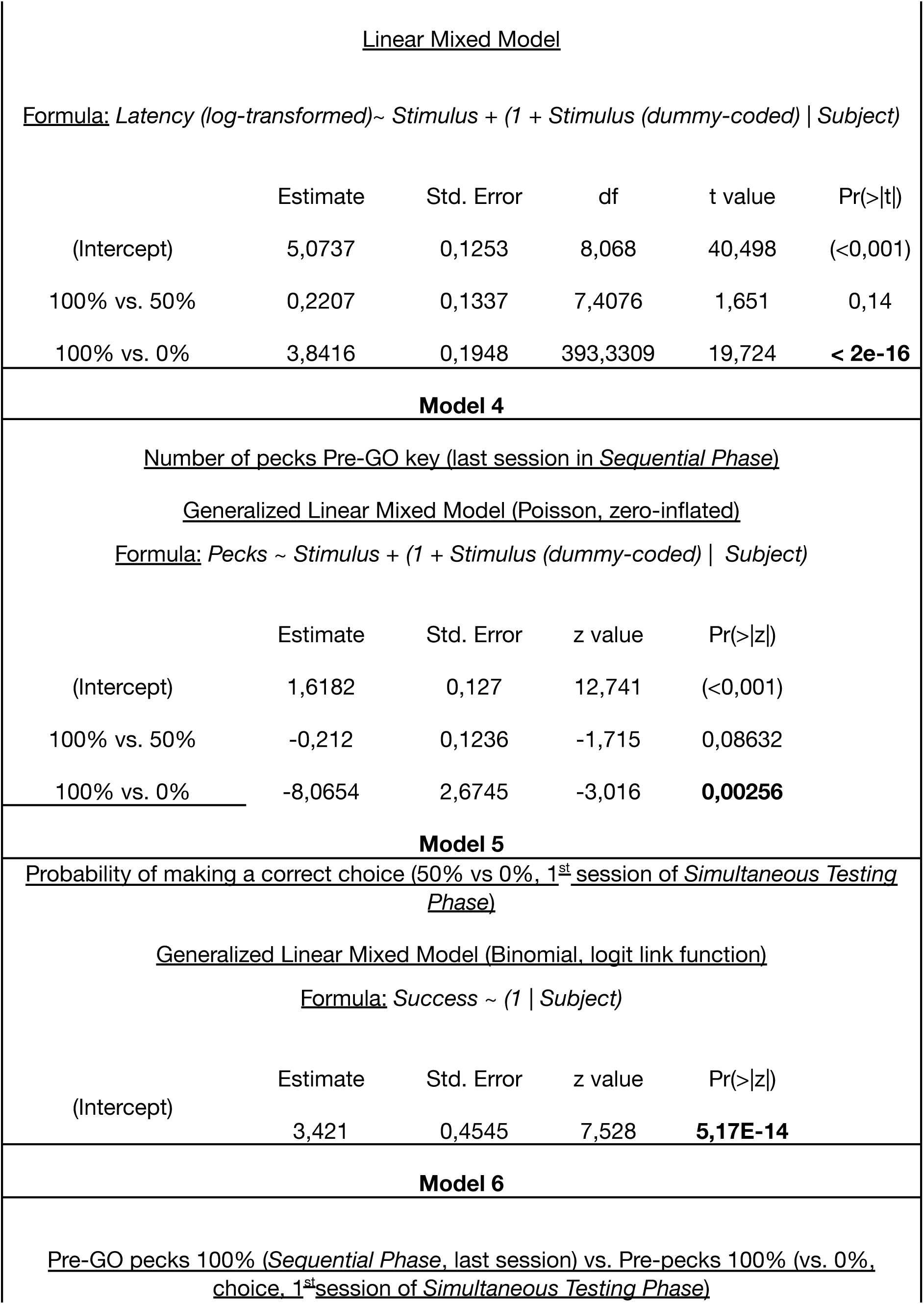
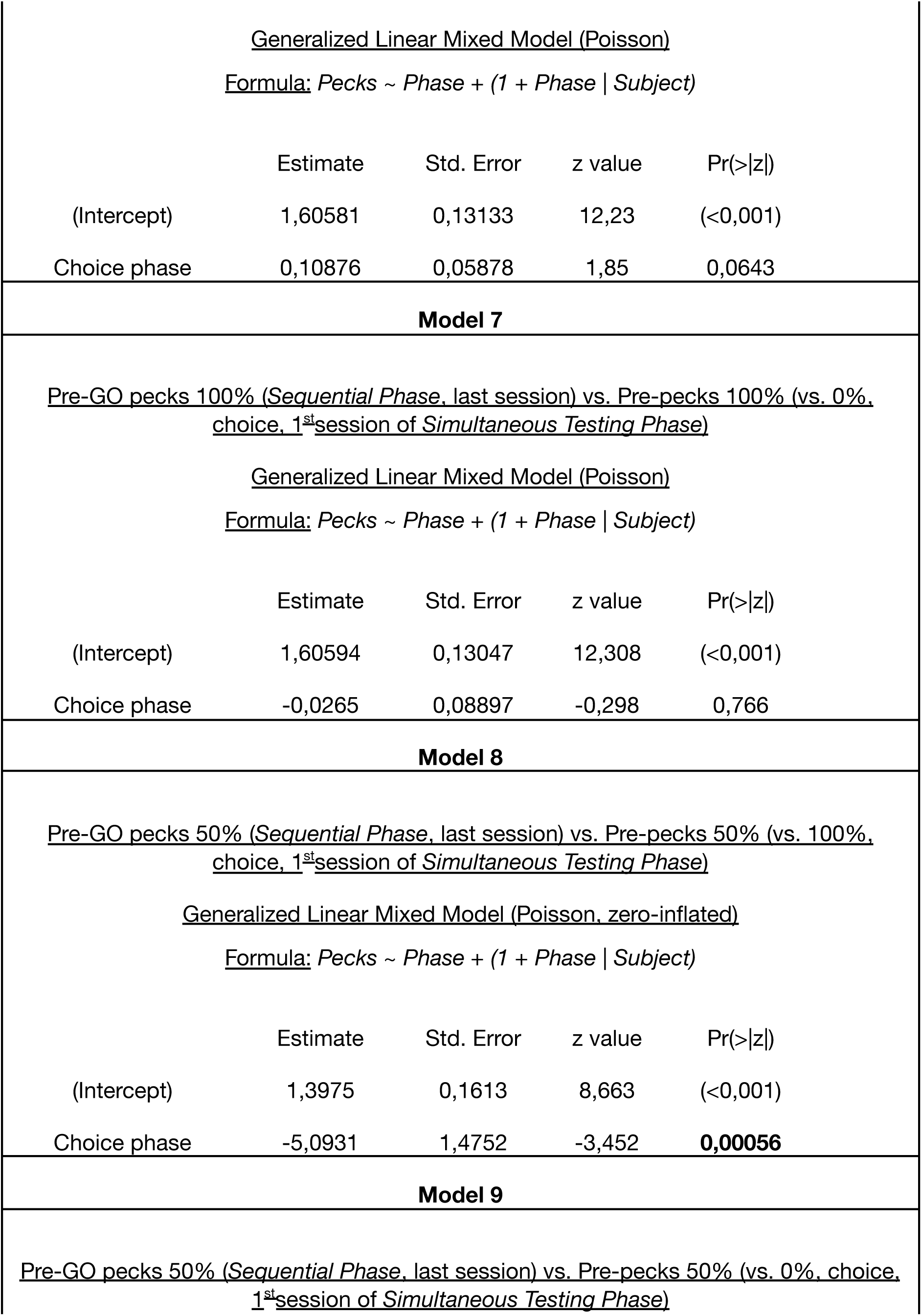
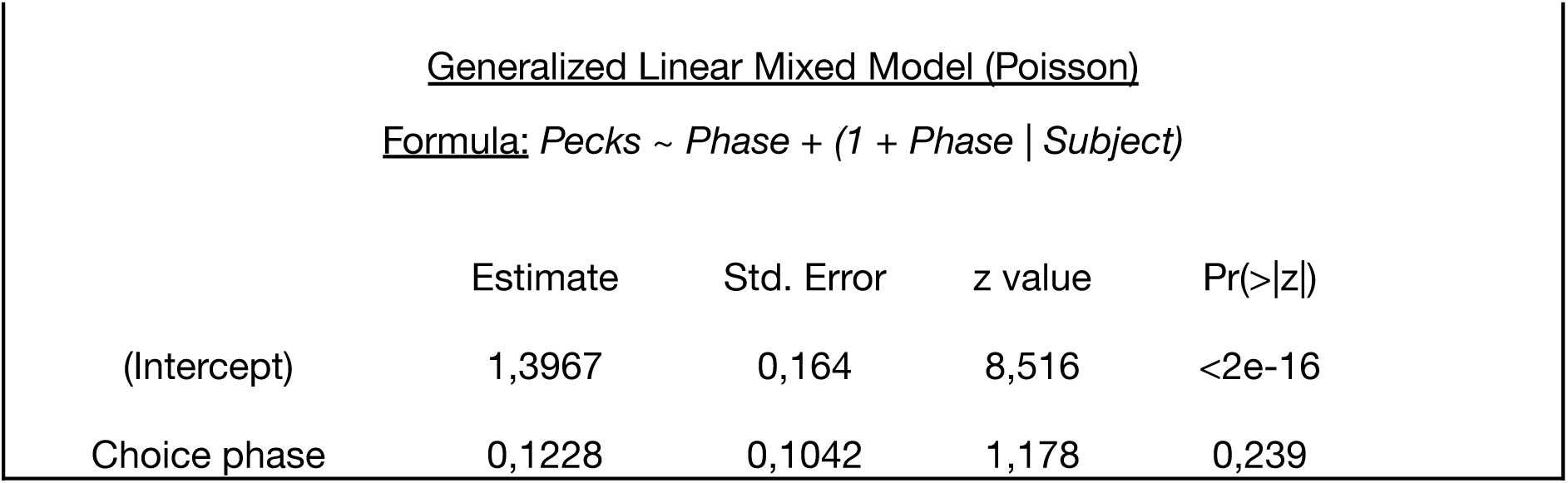
Statistical models formulations and output.

**Table S2.**
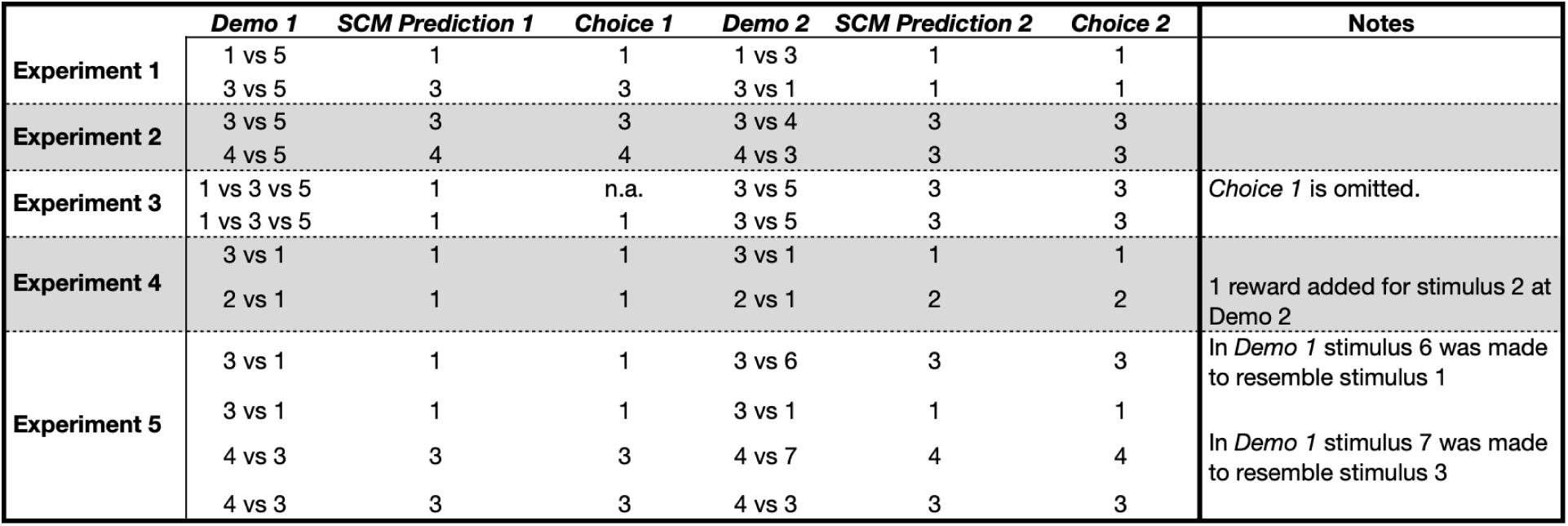
Schleihauf et al. (2025) performed five experiments with chimpanzees. They were inspired by the 2-Cups task, but each preference test included two, not one, sequences of a demonstration and a choice, as follows: *Demo 1 → Choice 1 → Demo 2 → Choice 2* This table summarizes Schleihauf et al. (2025) conditions, results and alternative predictions following a parallel processing of alternatives at the time of choice (Sequential Choice Model, SCM). Shleihauf et al used seven different event-stimuli, ranked qualitatively as candidates for ‘strength’ in association with reinforcement. They were: 1: food visible ≻ 2: food dropped, then invisible ≻ 3: cup shaken, food makes noise ≻ 4: food crumbs next to cup ≻ 5: cup undisturbed ≻ 6: food photo visible ≻ 7: stone visible The authors interpreted chimpanzees’ preferences in terms of initial (*Choice 1*) and revised (*Choice 2*) putative hypotheses. According to the SCM, we include predicted preferences based on utility and transitivity in all pairwise combinations (*SCM predictions 1* and *2*).

## Notes

### Competing Interest Statement

The authors have declared no competing interest.

### Summary of Updates

Title - updated for clarity; Abstract - shorter; Introduction and Discussion - Text updated and streamlined; Supplementary text and Table S2 added to Supplementary Materials.

