## Supplementary Materials for "Inferential reasoning in non-humans: a critical analysis of experimental evidence"

**The PDF file includes:**

Supplementary text
Figures S1 to S5
Table S1
Table S2

### Supplementary text

For the sequential training, the *Uncertain Reward* Pre-GO pecks decreased ($p = 0.031$) and response time increased ($p < 0.001$) when compared with the *Sure Reward* option (the model's intercept). However, no differences between stimuli were found at the last sequential training session (0.09 and 0.14 respectively). In order to make the informative options (*Sure Reward* and *No Reward*) more salient to the animals, we made it so that these two stimuli were blinking, while the *Uncertain Reward* stimulus was not. However, this made it so that, overall, the response key was always on for longer for the *Uncertain Reward* than for both the *Sure Reward* and *No Reward*. Given that the starlings had previous experience with the panels being active only when lit up, this may have caused them to press them less frequently (and therefore, have a lower response time on average as well) during the first few trials.

### Supplementary figures and tables

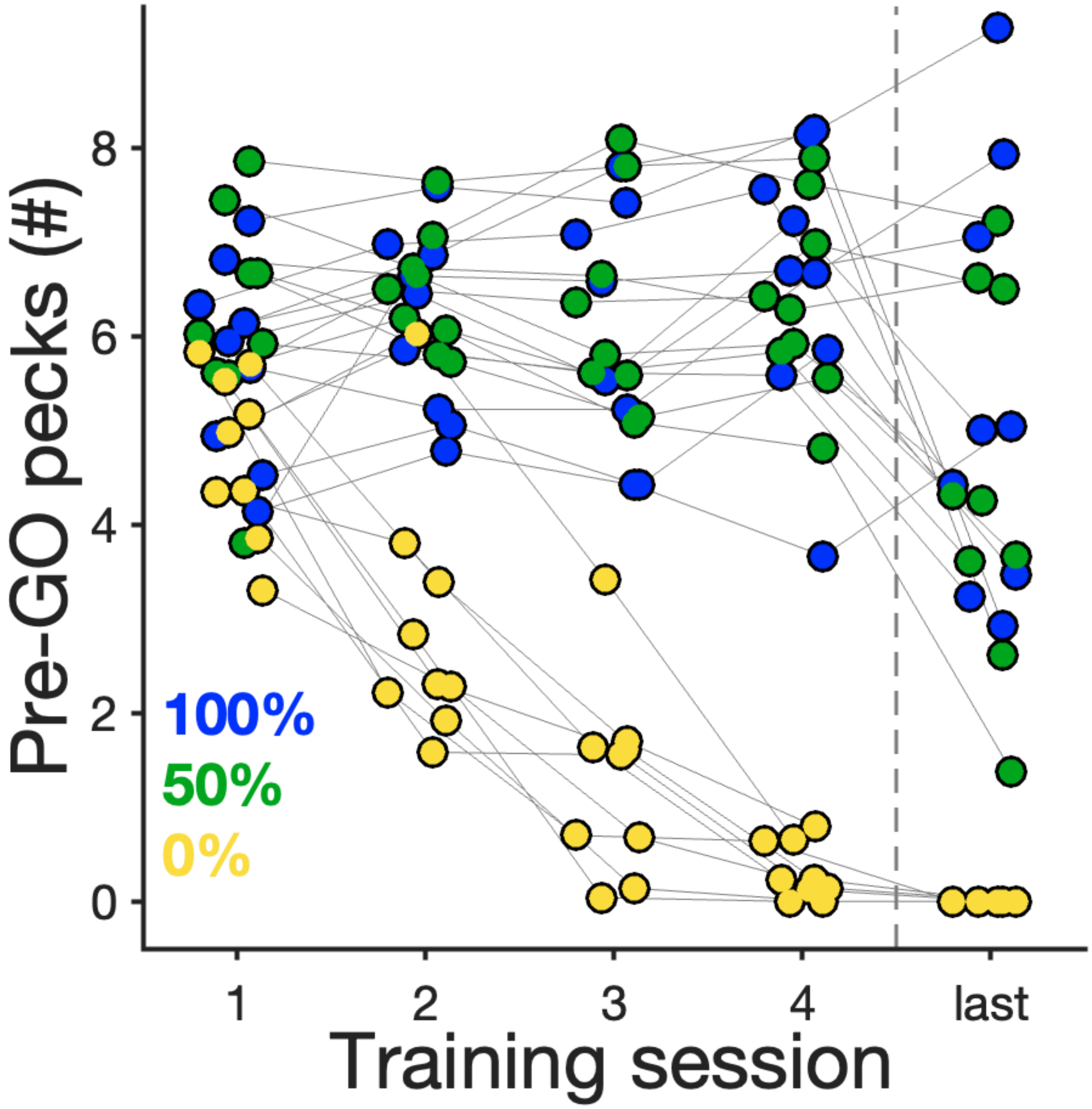


**Fig. S1.** Individual average Pre-GO pecks during the first 4 (and last) sessions of *Sequential Training*. Same data as presented in Figure 2A (bottom left).

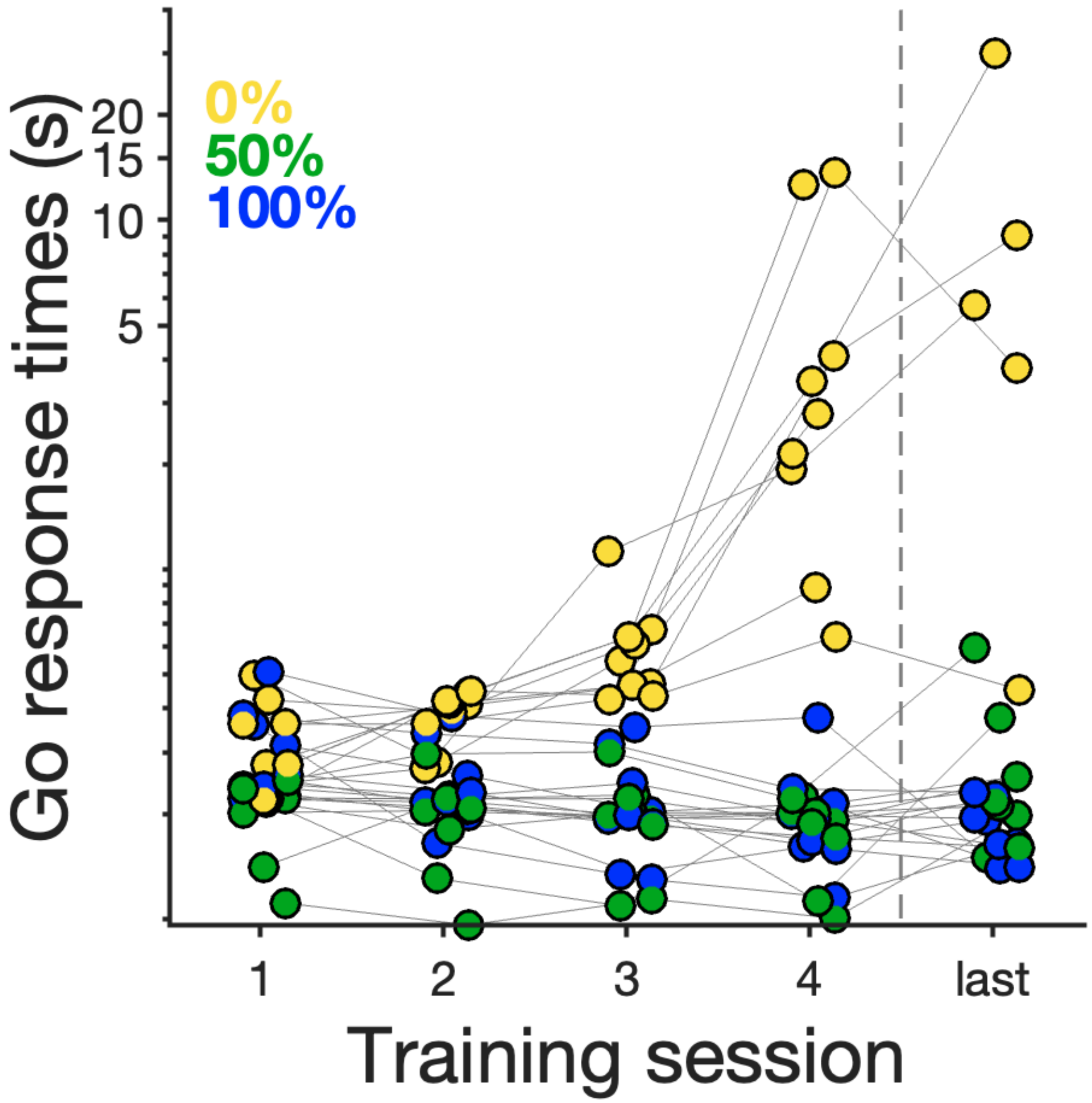


**Fig. S2.** Individual median response times to the Go cue during the first 4 (and last) sessions of *Sequential Training*. Same data as presented in Figure 2A (bottom right).

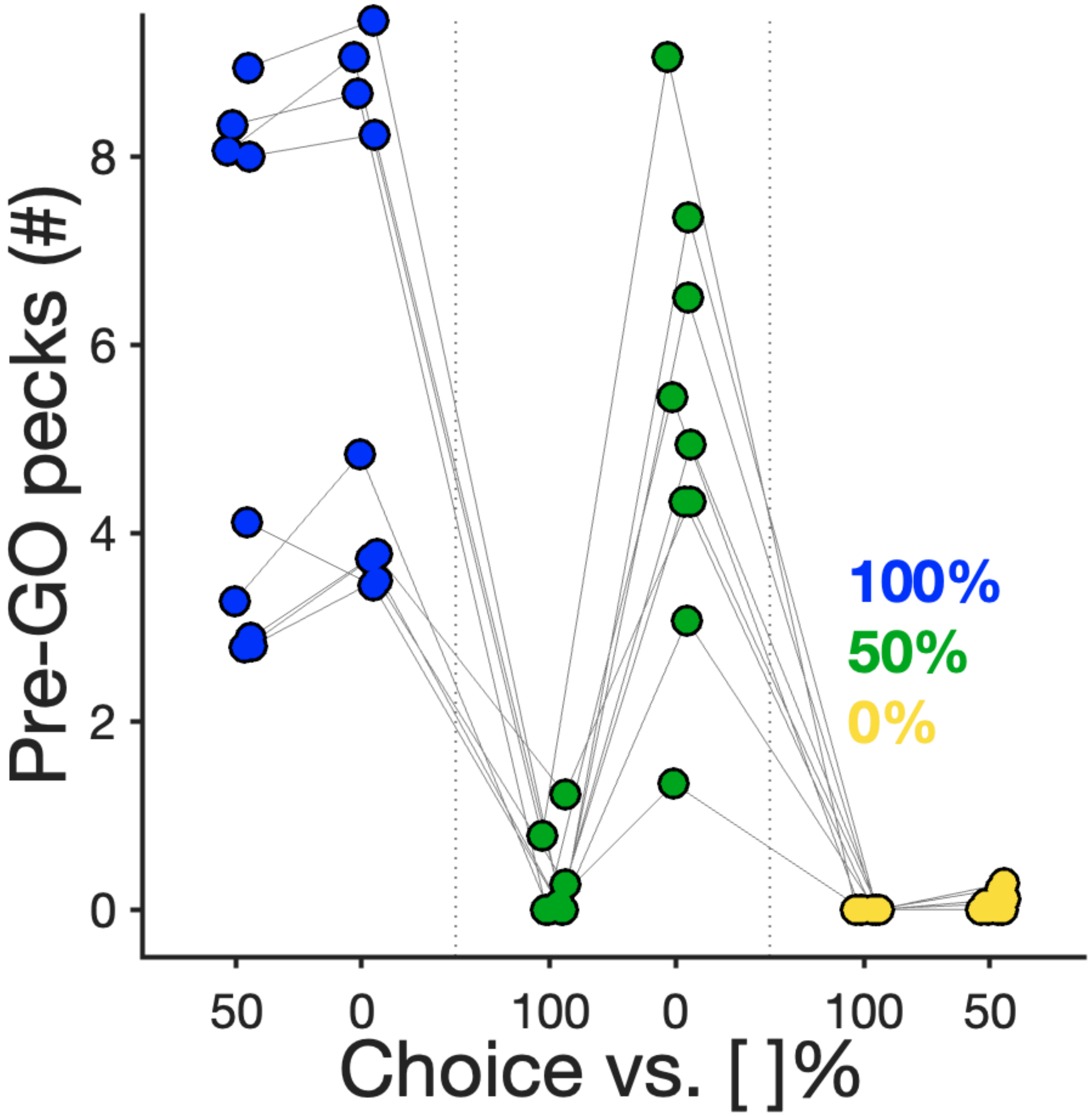


**Fig. S3.** Individual average Pre-GO pecks during the first session of *Simultaneous Testing*. Same data as presented in Figure 2B (bottom left).

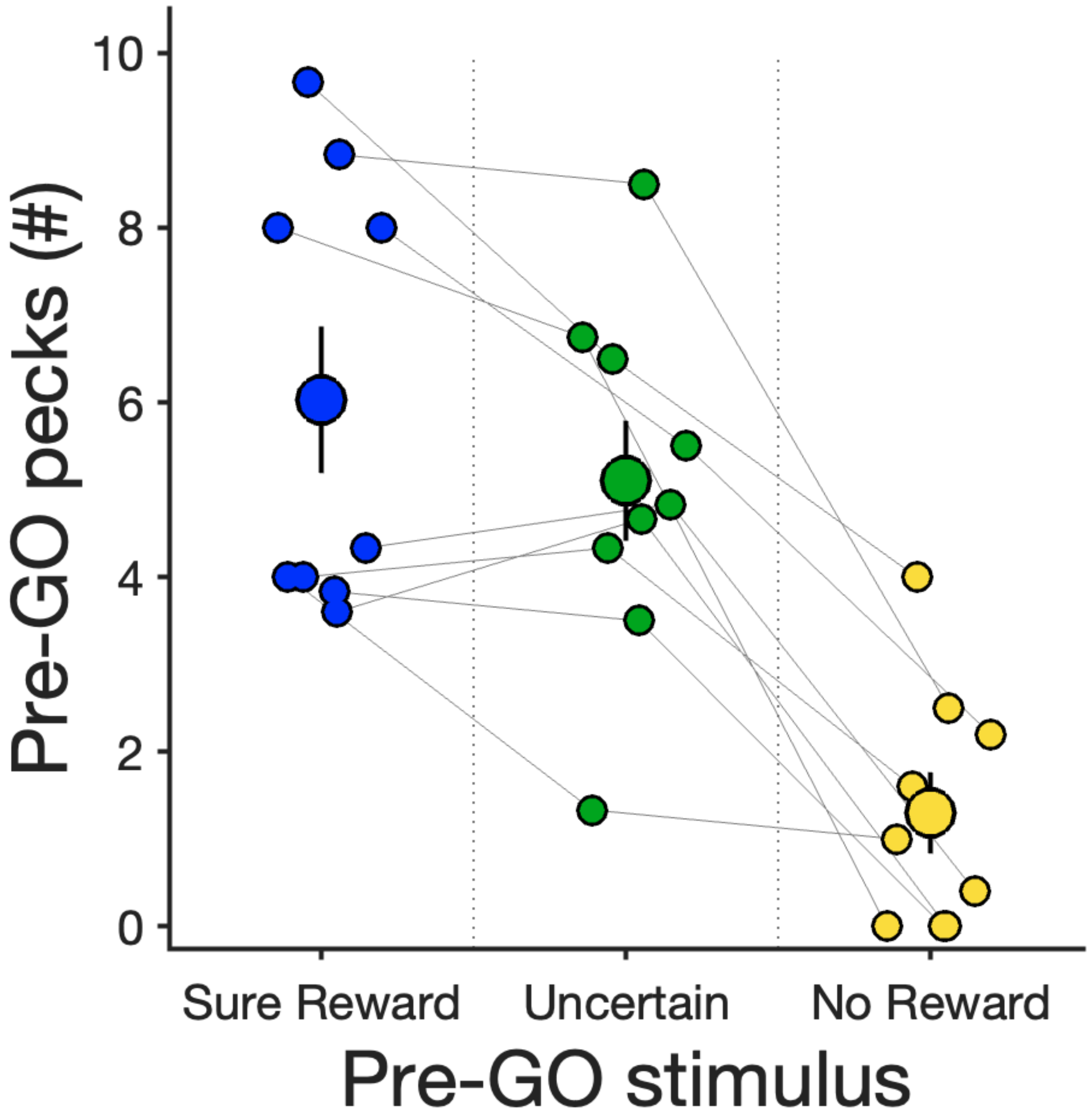


**Fig. S4.** Pre-GO pecks (mean±SE) for sequential trials during the first session of *Simultaneous Testing*. Smaller markers depict individual averages.

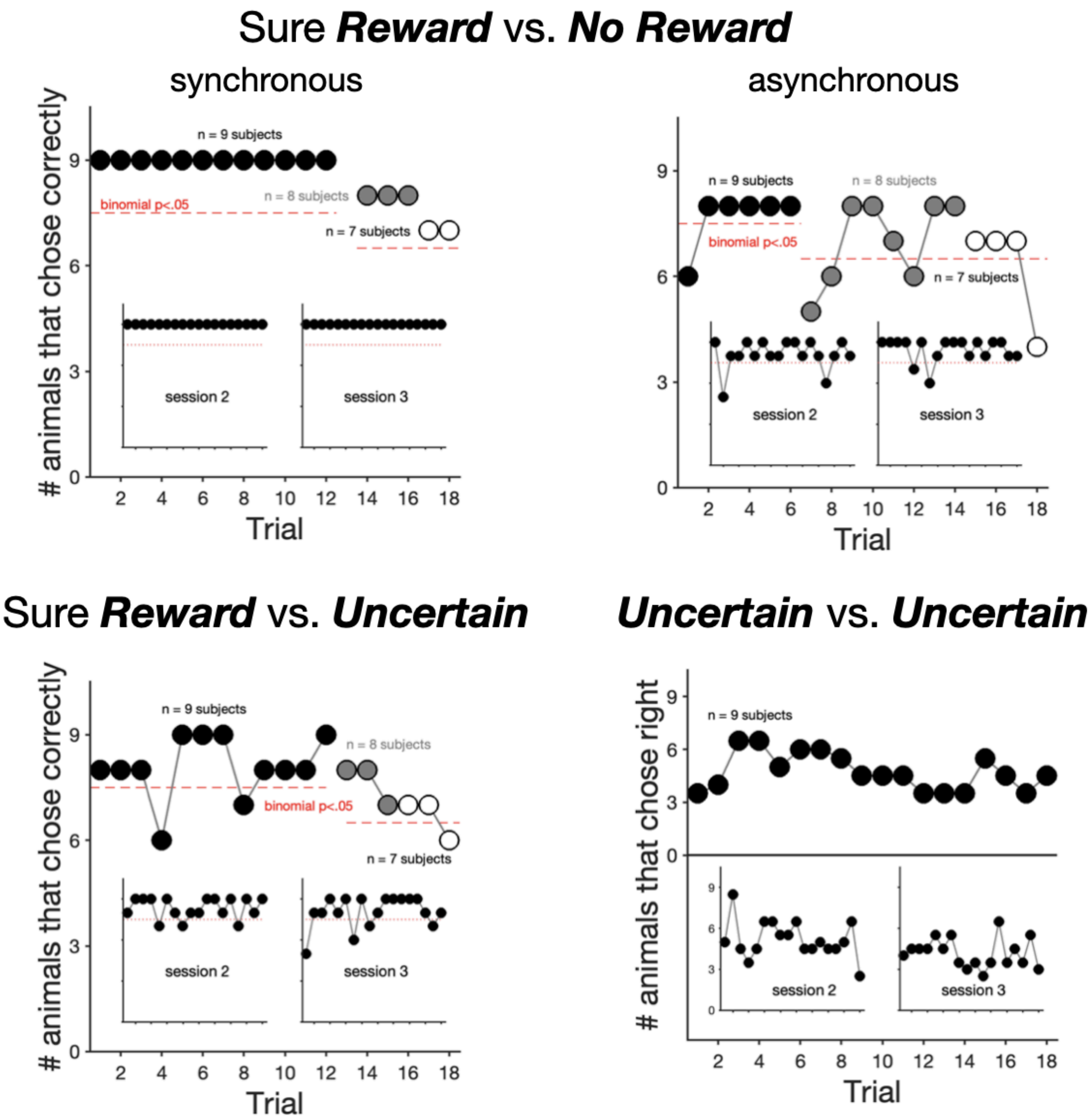


**Fig. S5.** Trial-by-trial number of animals responding correctly for the first session of *Simultaneous Testing* across the simultaneous choice trial types not shown in Figure 2B (bottom right). Marker colour depicts the number of subjects tested (black=9, grey=8 and white=7 subjects, respectively. Insets show the same data for testing sessions 2 and 3. Red dashed line depicts binomial significance. For *Uncertain* vs. *Uncertain* choice type, the y-axis represents the trial-by-trial number of animals that chose the right key, as there was no single correct (rewarded) response.

**Table S1.** Statistical models formulations and output.

| **Model 1.1** | | | | | |
|---|---|---|---|---|---|
| Latency to GO key (first four sessions in *Sequential Phase*) | | | | | |
| Linear Mixed Model | | | | | |
| Formula: *Latency (log-transformed) ~ Stimulus (dummy-coded) * Session (z-transformed) + (1 + Stimulus (dummy-coded) + Session (z-transformed) \| Subject)* | | | | | |
| | Estimate | Std. Error | df | t value | Pr(>\|t\|) |
| (Intercept) | 5,29623 | 0,11968 | 9,00105 | 44,253 | (<0,001) |
| 100% vs. 50% | -0,3452 | 0,10854 | 9,00405 | -3,18 | **0,01117** |
| 100% vs. 0% | 1,16712 | 0,1236 | 9,00986 | 9,443 | **5,71E-06** |
| Session (100%) | -0,1934 | 0,04701 | 15,29093 | -4,114 | **0,00089** |
| Session (50%) | 0,13821 | 0,03993 | 3724,172 | 3,461 | **0,00054** |
| Session (0%) | 1,00783 | 0,04199 | 3732,65 | 24,003 | **< 2e-16** |

| **Model 1.2** | | | | | |
|---|---|---|---|---|---|
| Latency to GO key (first four sessions in *Sequential Phase*) | | | | | |
| Linear Mixed Model | | | | | |
| Formula: *Latency (log-transformed) ~ Stimulus (dummy-coded) + Session (z-transformed) + (1 + Stimulus (dummy-coded) + Session (z-transformed) \| Subject)* | | | | | |
| | Estimate | Std. Error | df | t value | Pr(>\|t\|) |
| (Intercept) | 5,28223 | 0,11971 | 9,00006 | 44,126 | (<0,001) |
| 100% vs. 50% | -0,3396 | 0,10854 | 9,00105 | -3,128 | **0,01216** |

| | | | | | |
|---|---|---|---|---|---|
| 100% vs. 0% | 1,11495 | 0,12096 | 9,05086 | 9,217 | **6,75E-06** |
| Session | 0,14952 | 0,04024 | 9,01196 | 3,716 | **0,00479** |

**Model 2.1**

Number of pecks Pre-GO key (first four sessions in *Sequential Phase*)

Generalized Linear Mixed Model (Poisson, zero-inflated)

Formula: *Pecks ~ Stimulus (dummy-coded) * Session (z-transformed) + (1 + Stimulus (dummy-coded) + Session (z-transformed) | Subject)*

| | Estimate | Std. Error | z value | Pr(>\|z\|) |
|---|---|---|---|---|
| (Intercept) | 1,79365 | 0,06134 | 29,24 | (<0,001) |
| 100% vs. 50% | 0,04349 | 0,04594 | 0,95 | 0,3438 |
| 100% vs. 0% | -1,1978 | 0,02717 | -44,09 | **<2e-16** |
| Session (100%) | 0,03806 | 0,0241 | 1,58 | 0,1144 |
| Session (50%) | -0,0338 | 0,01569 | -2,16 | **0,0311** |
| Session (0%) | -0,8137 | 0,02581 | -31,52 | **<2e-16** |

**Model 2.2**

Number of pecks Pre-GO key (first four sessions in *Sequential Phase*)

Generalized Linear Mixed Model (Poisson, zero-inflated)

Formula: *Pecks ~ Stimulus (dummy-coded) + Session (z-transformed) + (1 + Stimulus (dummy-coded) + Session (z-transformed) | Subject)*

| | Estimate | Std. Error | z value | Pr(>\|z\|) |
|---|---|---|---|---|
| (Intercept) | 1,79729 | 0,06112 | 29,406 | (<0,001) |
| 100% vs. 50% | 0,04297 | 0,04347 | 0,989 | 0,3229 |
| 100% vs. 0% | -0,7346 | 0,03072 | -23,915 | **<2e-16** |
| Session | -0,0559 | 0,02354 | -2,374 | **0,0176** |

**Model 3**

Latency to GO key (last session in *Sequential Phase*)

Linear Mixed Model

Formula: *Latency (log-transformed)~ Stimulus + (1 + Stimulus (dummy-coded) | Subject)*

| | Estimate | Std. Error | df | t value | Pr(>\|t\|) |
|---|---|---|---|---|---|
| (Intercept) | 5,0737 | 0,1253 | 8,068 | 40,498 | (<0,001) |
| 100% vs. 50% | 0,2207 | 0,1337 | 7,4076 | 1,651 | 0,14 |
| 100% vs. 0% | 3,8416 | 0,1948 | 393,3309 | 19,724 | **< 2e-16** |

**Model 4**

Number of pecks Pre-GO key (last session in *Sequential Phase*)

Generalized Linear Mixed Model (Poisson, zero-inflated)

Formula: *Pecks ~ Stimulus + (1 + Stimulus (dummy-coded) | Subject)*

| | Estimate | Std. Error | z value | Pr(>\|z\|) |
|---|---|---|---|---|
| (Intercept) | 1,6182 | 0,127 | 12,741 | (<0,001) |
| 100% vs. 50% | -0,212 | 0,1236 | -1,715 | 0,08632 |
| 100% vs. 0% | -8,0654 | 2,6745 | -3,016 | **0,00256** |

**Model 5**

Probability of making a correct choice (50% vs 0%, 1st session of *Simultaneous Testing Phase*)

Generalized Linear Mixed Model (Binomial, logit link function)

Formula: *Success ~ (1 | Subject)*

| | Estimate | Std. Error | z value | Pr(>\|z\|) |
|---|---|---|---|---|
| (Intercept) | 3,421 | 0,4545 | 7,528 | **5,17E-14** |

**Model 6**

Pre-GO pecks 100% (*Sequential Phase*, last session) vs. Pre-pecks 100% (vs. 0%, choice, 1st session of *Simultaneous Testing Phase*)

Generalized Linear Mixed Model (Poisson)

Formula: *Pecks ~ Phase + (1 + Phase | Subject)*

| | Estimate | Std. Error | z value | Pr(>\|z\|) |
|---|---|---|---|---|
| (Intercept) | 1,60581 | 0,13133 | 12,23 | (<0,001) |
| Choice phase | 0,10876 | 0,05878 | 1,85 | 0,0643 |

**Model 7**

Pre-GO pecks 100% (*Sequential Phase*, last session) vs. Pre-pecks 100% (vs. 0%, choice, 1st session of *Simultaneous Testing Phase*)

Generalized Linear Mixed Model (Poisson)

Formula: *Pecks ~ Phase + (1 + Phase | Subject)*

| | Estimate | Std. Error | z value | Pr(>\|z\|) |
|---|---|---|---|---|
| (Intercept) | 1,60594 | 0,13047 | 12,308 | (<0,001) |
| Choice phase | -0,0265 | 0,08897 | -0,298 | 0,766 |

**Model 8**

Pre-GO pecks 50% (*Sequential Phase*, last session) vs. Pre-pecks 50% (vs. 100%, choice, 1st session of *Simultaneous Testing Phase*)

Generalized Linear Mixed Model (Poisson, zero-inflated)

Formula: *Pecks ~ Phase + (1 + Phase | Subject)*

| | Estimate | Std. Error | z value | Pr(>\|z\|) |
|---|---|---|---|---|
| (Intercept) | 1,3975 | 0,1613 | 8,663 | (<0,001) |
| Choice phase | -5,0931 | 1,4752 | -3,452 | **0,00056** |

**Model 9**

Pre-GO pecks 50% (*Sequential Phase*, last session) vs. Pre-pecks 50% (vs. 0%, choice, 1st session of *Simultaneous Testing Phase*)

Generalized Linear Mixed Model (Poisson)

Formula: *Pecks ~ Phase + (1 + Phase | Subject)*

| | Estimate | Std. Error | z value | Pr(>\|z\|) |
|---|---|---|---|---|
| (Intercept) | 1,3967 | 0,164 | 8,516 | <2e-16 |
| Choice phase | 0,1228 | 0,1042 | 1,178 | 0,239 |

**Table S2.** Schleihauf et al. (2025) performed five experiments with chimpanzees. They were inspired by the 2-Cups task, but each preference test included two, not one, sequences of a demonstration and a choice, as follows:

*Demo 1 → Choice 1 → Demo 2 → Choice 2*

This table summarizes Schleihauf et al. (2025) conditions, results and alternative predictions following a parallel processing of alternatives at the time of choice (Sequential Choice Model, SCM). Shleihauf et al used seven different event-stimuli, ranked qualitatively as candidates for 'strength' in association with reinforcement. They were:

1: food visible > 2: food dropped, then invisible > 3: cup shaken, food makes noise > 4: food crumbs next to cup > 5: cup undisturbed > 6: food photo visible > 7: stone visible

The authors interpreted chimpanzees' preferences in terms of initial (*Choice 1*) and revised (*Choice 2*) putative hypotheses. According to the SCM, we include predicted preferences based on utility and transitivity in all pairwise combinations (*SCM predictions 1* and *2*).

| | *Demo 1* | *SCM Prediction 1* | *Choice 1* | *Demo 2* | *SCM Prediction 2* | *Choice 2* | Notes |
|---|---|---|---|---|---|---|---|
| **Experiment 1** | 1 vs 5 | 1 | 1 | 1 vs 3 | 1 | 1 | |
| | 3 vs 5 | 3 | 3 | 3 vs 1 | 1 | 1 | |
| **Experiment 2** | 3 vs 5 | 3 | 3 | 3 vs 4 | 3 | 3 | |
| | 4 vs 5 | 4 | 4 | 4 vs 3 | 3 | 3 | |
| **Experiment 3** | 1 vs 3 vs 5 | 1 | n.a. | 3 vs 5 | 3 | 3 | *Choice 1* is omitted. |
| | 1 vs 3 vs 5 | 1 | 1 | 3 vs 5 | 3 | 3 | |
| **Experiment 4** | 3 vs 1 | 1 | 1 | 3 vs 1 | 1 | 1 | |
| | 2 vs 1 | 1 | 1 | 2 vs 1 | 2 | 2 | 1 reward added for stimulus 2 at Demo 2 |
| **Experiment 5** | 3 vs 1 | 1 | 1 | 3 vs 6 | 3 | 3 | In *Demo 1* stimulus 6 was made to resemble stimulus 1 |
| | 3 vs 1 | 1 | 1 | 3 vs 1 | 1 | 1 | |
| | 4 vs 3 | 3 | 3 | 4 vs 7 | 4 | 4 | In *Demo 1* stimulus 7 was made to resemble stimulus 3 |
| | 4 vs 3 | 3 | 3 | 4 vs 3 | 3 | 3 | |
